# HIF-1 activated by PIM1 assembles a pathological transcription complex and regulon that drives JAK2V617F MPN disease

**DOI:** 10.1101/2024.07.02.601518

**Authors:** David Kealy, Ruth Ellerington, Suraj Bansal, Andy G.X. Zeng, Jessie J.F. Medeiros, Katie A. West, Nicole-Mae Blacknell, Catherine A. Hawley, Jakub Lukaszonek, Richard T. Gawne, Hwei Minn Khoo, Gillian Caalim, Bryce Drylie, Jenny Chatzigerou, Bianca Lima Ferreira, Adele K Fielding, Guanlin Wang, David G. Kent, Bethan Psaila, Adam C. Wilkinson, Andrew N. Holding, Ian S. Hitchcock, Andrew S. Mason, Vikas Gupta, John E. Dick, Katherine S. Bridge

**Affiliations:** Centre for Blood Research, University of York; York, United Kingdom; York Biomedical Research Institute, University of York; York, United Kingdom; Princess Margaret Cancer Centre, University Health Network; Toronto, Canada; University of Toronto; Toronto, Canada; Barts Cancer Institute, Queen Mary University of London; London, UK; Jack Birch Unit for Molecular Carcinogenesis, University of York; York, UK; Hull York Medical School; Medical Research Council Weatherall Institute of Molecular Medicine (MRC WIMM) and NIHR Biomedical Research Centre Haematology Theme; University of Oxford; Oxford, UK; MRC WIMM Centre for Computational Biology, University of Oxford; Oxford, United Kingdom; Shanghai Key Laboratory of Metabolic Remodeling and Health, Institute of Metabolism and Integrative Biology; Fudan University, Shanghai, China; Qizhi Institute, Shanghai, China; Oxford University Hospitals NHS Trust; Oxford, UK

**Author notes:** These authors contributed equally (David Kealy, Ruth Ellerington). **Data Sharing** Data for this study are available at NCBI SRA PRJNA1144472.

## Abstract

Hypoxia-inducible factors (HIFs) are master transcriptional regulators, central to cellular survival under limited oxygen (hypoxia) and frequently activated within malignancy. Malignant context affects the role of HIFs within oncogenesis; however, the mechanisms regulating HIF context-specificities are not well characterised. Applying the JAK2V617F (JVF) model of myeloproliferative neoplasms (MPNs), in which HIF-1 is active in normoxia (20% O_2_), we sought to determine whether the modality of HIF-1 activation directs its function. We identify that HIF-1 is stabilised in JVF cells downstream of STAT1/5 signalling and upregulation of PIM1: PIM1 mediates phosphorylation of HIF-1 (Thr498/Ser500) in JVF cells that inhibits proteasomal degradation. PIM1 inhibition eradicates HIF-1 from JVF cells. Applying a single-input dual-omics output chromatin interactome methodology (DOCIA), we define JVF-specific transcription cofactors and genomic redistribution of HIF-1, and a JVF-HIF-1 regulon in primary haematopoietic stem/progenitor cells. In a cohort of 172 JVF-MPN patients, we observe significant association of the JVF-HIF-1 regulon (but strikingly, not canonical HIF-1 genes) with disease severity, progression, and patient survival. Finally, we identify a core set of JVF-HIF-1 targets significantly associated with spontaneous transformation of MPNs to AML. Our findings identify that HIF-1 activation by the JVF-PIM1 axis substantially alters its function, and that this reprogramming drives MPN disease progression, restoring the potential for targeted therapies that delineate HIF-1 activity co-opted by malignancy from essential roles within physiological oxygen homeostasis.

**Key Points:**

1. HIF-1 activation via PIM1 in JAK2V617F-MPNs drives non-canonical transcription complex formation/function.
2. The JAK2V617F-HIF-1 regulon drives MPN disease progression, transformation to AML and worse patient outcomes.

## Introduction

Hypoxia-inducible factors (HIFs) are the major transcriptional regulators of cellular oxygen homeostasis. Under normal oxygen conditions, prolyl hydroxylase domain proteins (PHDs) hydroxylate two highly conserved residues on HIF-1α (Pro402/Pro564), resulting in its recognition by the von Hippel-Lindau (VHL) E3 ligase complex, ubiquitination, and degradation by the proteasome^1^. During hypoxia, limited oxygen inhibits PHD activity, and stabilised HIF-1α protein translocates to the nucleus; the resulting heterodimerisation with the constitutively expressed HIF-1β/ARNT^2^ and ensuing recruitment of transcription coactivators forms an active transcription complex that binds to hypoxia-response elements (HREs) within target gene promoters^3^, initiating the activation of a wide array of genes involved in the response to hypoxia.

Whilst critical to the physiological response to hypoxic exposure of normal cells, the portfolio of genes regulated by HIF provide a competitive advantage to cancer cells. In solid and haematological malignancies, HIFs are overexpressed as a result of two major factors: by hypoxic conditions - such as tumour or bone marrow (BM) microenvironments- and by genetic mutations, including *VHL*, *p53*, *Bcl2*, *Myc*, *Ras* and *JAK*^1^. In contrast to HIF-2, which is exclusively oncogenic, HIF-1 confers either oncogenic or tumour suppressive functions^2^. In myeloid neoplasia, inhibition of HIF-1 is efficacious for treatment of MPNs^4^, whereas HIF-1 activation was recently evidenced for treatment of acute myeloid leukaemia (AML)^5^. The underpinning mechanisms that orchestrate the multiplicity of HIF-1 within malignancy are poorly understood.

To reconcile the evident context-specificity of HIF-1 activity within malignant haematopoiesis, we sought to determine whether the mechanism of HIF-1 activation determines its function. The JAK2V617F (JVF) point mutation is highly prevalent within philadelphia chromosome negative MPNs and is carried by >95% of patients with polycythaemia vera (PV), 50-60% of patients with essential thrombocythemia (ET) and primary myelofibrosis (PMF)^6–8^. As HIF-1α is stabilised in JVF-mutant MPNs in normoxic conditions^9^, we applied this model to determine whether the mutational context/modality of HIF-1 activation determines its function. Herein we demonstrate that PIM1-mediated stabilisation of HIF-1α downstream of JVF signalling significantly alters the formation and function of HIF-1 transcription complexes compared to hypoxic activation, enacting a JVF-specific HIF-1 regulon that is significantly associated with disease progression and survival of JVF MPN patients.

## Methods

### Dual-Output Chromatin Interactome Assay (DOCIA)

BaF3 hMPL hJAK2 WT or BaF3 hMPL hJAK2 VF cells were cultured in indicated conditions, then fixed, pelleted by centrifugation, resuspended into formaldehyde and neutralised by the addition of glycine. Sample processing and HIF-1α chromatin-immunoprecipitations were performed as described previously^10,11^. Samples were split and processed through either next generation sequencing or mass spectrometry to identify HIF-1α binding loci or transcription cofactors respectively. Comprehensive DOCIA method details can be found along with full study methods in Supplementary Information.

## Results

### HIF-1α is stabilised in JAK2V617F cells downstream of disproportionately activated STAT1/STAT5 signalling and increased PIM1 kinase expression

First, we confirmed that HIF-1α stabilisation^9^ occurred throughout a panel of JVF mutant cell-line models (BaF3/hMPL^12,13^, UT7/TPO ^13,14^, SET2^15,16^ and HEL^17^) (Fig.1A). HIF-1α protein levels in Lin^-^jak2^wt/wt^, jak2^wt/v617f^ and jak2^v617f/v617f^ C57/BL6 mouse BM^18^ also correlated with JVF allelic-burden (Fig.1B;Fig.S1A-C). We previously demonstrated that hypoxia enhances the purity of murine HSCs in *ex vivo* PVA expansion culture^19^; applying this system we confirmed that LT-HSCs express oxygen-correlative levels of HIF-1α protein (Fig.1C), upon which they are dependent for expansion (GN44028; HIF-1 inhibitor [HIF-1i])^20,21^ (Fig.1D; Fig.S2). JVF hom LT-HSCs expansion was not supported by this system, however JVF het LT-HSCs grew in culture correspondingly with the immunophenotypic ratios from fresh BM ^18^, (Fig.S3A-B). HIF-1i suppressed the LSK fraction irrespective of phenotype, although JVF-specificity was observed in reducing Prog expansion. These data demonstrated an essential role of HIF-1 within WT and JVF-HSPCs, thereby compelling the delineation of HIF-1 activation in each of these contexts.

**Figure 1.**
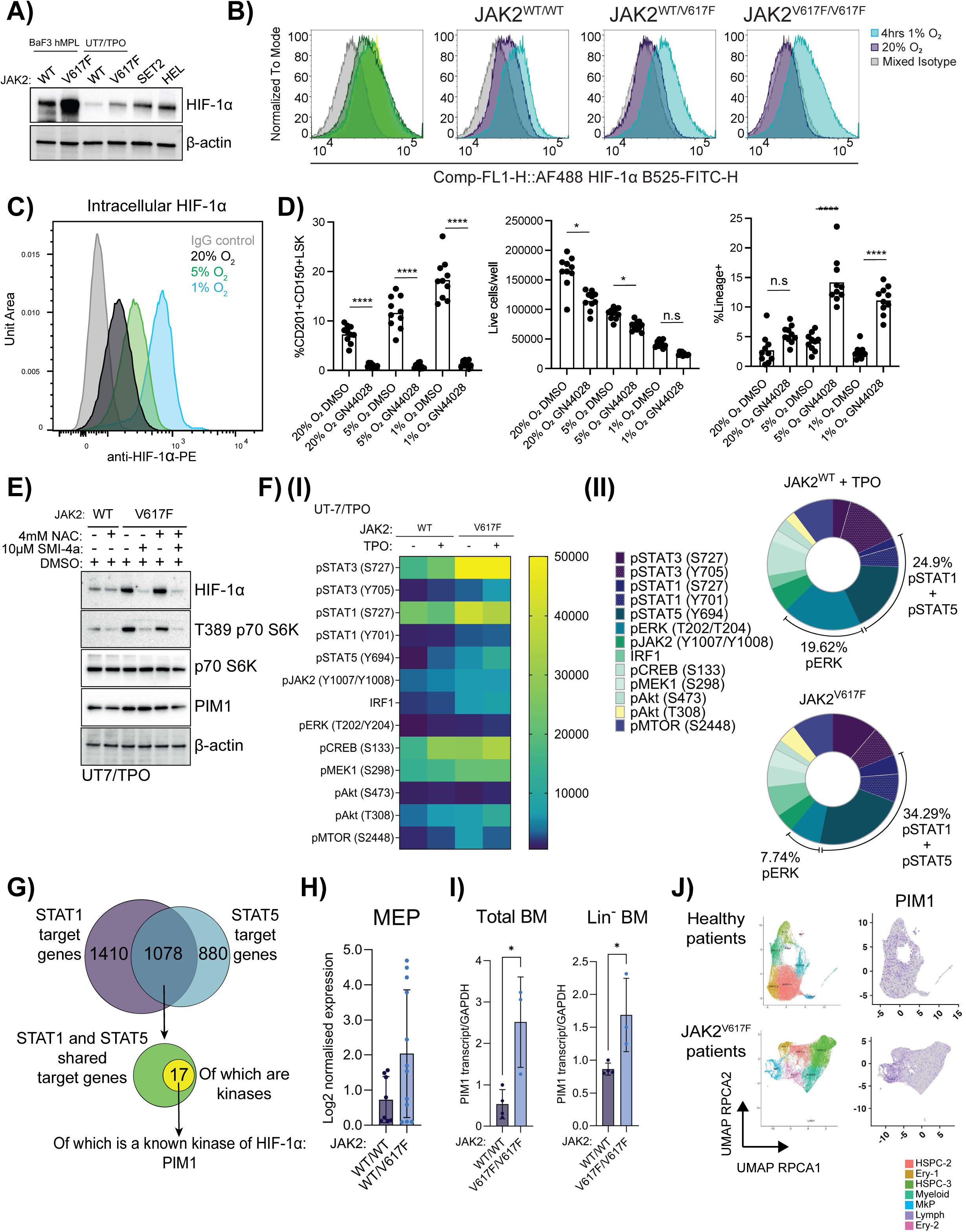
(A) JVF stabilises HIF-1α independent of oxygen levels in monogenic JVF and JVF MPN immortalised cell lines. Immunoblot of HIF-1α levels in normoxic cultured wildtype and JVF-monogenic UT7/TPO and hMPL BaF3 cells, JVF heterozygous MPN-patient derived SET2 cells and JVF homozygous MPN-patient derived HEL cells. Immunoblotting carried out using HIF-1α antibody with β actin as a loading control (n=3). (B) JVF stabilises HIF-1α in primary mouse lineage depleted BM cells. Flow cytometry histogram distribution of channel fluorescent intensity (normalised to mode) and cell count (table below) from JVF homozygous (n=1), heterozygous (n=2) and wildtype (n=2) C57/BL6 mouse lineage-depleted BM cells probed with fluorescently labelled HIF-1α antibody, following ex-vivo culture in 4hrs hypoxic 1% O2 (blue) or normoxic 20% O2 (purple) conditions (n=2). (C) In wildtype primary mouse HSCs, HIF-1α is stabilised by physioxic (5% O2) and hypoxic (1% O2) conditions correlative to oxygen levels. Representative intracellular flow cytometry plot for anti-HIF-1α -PE staining in wildtype mouse HSC-derived 20%, 5%, or 1% O2 cultures. (D) Treatment with HIF-1α inhibitor reduces the population of CD201+CD150+LSK cells and overall cell viability whilst increasing the percentage of lineage committed cells in 10-day PVA expansion culture. Mean frequency of CD201+CD150+LSK cells (left), mean number of live cells per well (middle), and mean frequency of Lineage+ cells (right) within 10-day HSC-derived cultures at 20% O2, 5% O2, or 1% O2, cultured with either DMSO or the HIF-1ɑ-specific transcription inhibitor GN44028 (100 nM) (n=10). (E) Treatment with ROS scavenger NAC (N-acetyl-L-cysteine) fails to inhibit JVF-induced HIF-1α stabilisation (whereas PIM1 inhibition with SMI-4a, does inhibit JVF-induced HIF-1α stabilisation). Immunoblot of HIF-1α, PIM1 and PIM1 substrate (p70 S6K) levels in wildtype and JVF UT7/TPO cells starved overnight and treated as indicated with either NAC (from the onset of starvation - 24hr treatment), SMI-4a (6hr treatment) or vehicle control. Immunoblotting carried out using HIF-1α, T389 p70 S6K, p70 S6K and PIM1 antibodies with β actin as a loading control (n=3). (F)(I) Kinase cascade activity downstream of JAK2 shows significant differences between TPO-stimulated JAK2 and JVF-mediated constitutive signalling. Wildtype and JVF UT7/TPO cells overnight starved or treated with TPO (thrombopoietin) and analysed by multiplex fluorescent phospho-flow cytometry. Heatmap of mean fold change of channel fluorescence intensity (MFI), data normalised based on fold change to mean value of wildtype starved, no TPO, normoxic condition (±SEM) (n=3). (II) Proportional representation of kinase activity as detected by phospho-flow cytometry relative to total kinase cascade signalling downstream of JAK2 shows an increase in the relative amount of STAT1 and STAT5 activation in JVF cells compared to TPO/MPL stimulation of wildtype JAK2 (n=3). (G) Database analysis of STAT1 and STAT5 target genes show PIM1 as the sole known kinase of HIF-1α. STAT1 gene targets indicated in purple, STAT5 gene targets indicated in blue, shared STAT1 and STAT5 targets indicated in green, of which are kinases shown in yellow. (H) PIM1 transcript expression is significantly higher in JVF mouse MEPs (Megakaryocytic-erythroid progenitors) compared to wildtype. Bulk RNA-seq analysis of mouse MEPs from wildtype (n=3) and JVF heterozygous mice (n=3). (I) PIM1 transcript expression is significantly higher in both total and lineage negative JVF mouse BM. qRT-PCR of PIM1 transcript levels in homozygous JVF (n=3) and wildtype (n=3) mouse BM HSPCs (total and lineage negative) with GAPDH as a housekeeping gene control. (J) PIM1 transcript levels are elevated and localised to the megakaryocyte progenitor (MkP) population in JVF positive MPN patients when compared to healthy controls. UMAP scRNAseq plots of control (top row) and myelofibrosis (bottom row) HSPCs with PIM1 transcript highlighted in purple.

Whilst previous findings implicated reactive oxygen species (ROS) as stabilising HIF-1α downstream of TPO/JVF signalling^9,22^, application of a ROS scavenger (NAC) did not destabilise HIF-1α in the JVF cell-line panel (Fig.1E;Fig.S4A-B). Phosphorylation of HIF-1 critically modulates its interactome/function^23–28^; we therefore reasoned that kinase activation downstream of JVF could be the responsible mechanism for HIF-1α stabilisation. ERK1/2 is an essential HIF-1 kinase^29^, and is hyperactivated downstream of JVF mutation^30^, however ERK1/2 specific inhibitors also failed to destabilise HIF-1α protein in JVF cells (Fig.S5). We therefore applied an unbiased, high-throughput phospho-flow cytometry approach (Fig.S6)^31^ to fully characterise the signalling cascades downstream of JVF mutation in UT-7/TPO WT/JVF cells. We observed not only global amplification of signalling in JVF mutant cells vs. TPO stimulation but identified disproportionate activation of specific downstream kinase cascades (Fig.1F[I]); STAT1/STAT5 were hyperactivated relative to other cascades, collectively representing 34.3% of downstream signalling output in JVF cells (vs. 24.9%; TPO-stimulated WT JAK2 cells) (Fig.1F[II]). This was coupled with reduced relative ERK activation, supporting our earlier findings excluding ERK-mediated HIF-1α phosphorylation (Fig.S5). Unlike STAT3, STAT1/5 are not known to be direct protein-protein interactors of HIFs^32,33^; we therefore interrogated STAT1/5 target genes for known kinase regulators of HIF-1α^34–37^, identifying 1078common gene targets between STAT1/STAT5 (Fig.1G), within which 17 kinases were identified. Of these, PIM1 is a known HIF-1α-kinase^38^, is upregulated in murine JVF HSPCs^30^, and contributes to the pathogenesis of haematological malignancies^7^. We identified upregulation of PIM1 transcript in LSK/megakaryocyte erythroid progenitor (MEP)s in JVF het mice^39^ (Fig.1H;Fig.S7), which we confirmed via qRT-PCR of whole/Lin-BM (Fig.1I). Analysis of scRNAseq data^40^ for JVF MPN patients, demonstrated elevated total PIM1 and overexpression within the disease-driving expanded megakaryocyte progenitor (MkP) population (Fig.1J).

### PIM1 inhibition confirms JAK2V617F-induced PIM1 stabilises HIF-1α without modifying JVF signalling and does so via phosphorylation of Thr498/Ser500

Application of PIM1-specific inhibitors (PIMi) (TCS PIM1; SMI-4a) and a pan-PIM inhibitor (PIM447) to WT/JVF isogenic cell lines revealed dose-dependent decreases of normoxic HIF-1α protein in JVF cells with all PIMi (Fig.2A(I); Fig.S8-9). Analysis of signalling cascades by phospho-/flow in UT7/TPO WT/JVF cells treated with either SMI-4a (PIM1i) or Ruxolitinib (Rux)^41–45^ under a range of conditions revealed that SMI-4a reduced HIF-1α levels in JVF cells in both normoxia and hypoxia, however had no effect on HIF-1α levels in WT cells stimulated with TPO in either oxygen condition (Fig.2A[II]). These findings demonstrate that inhibition of PIM1 can confer specificity to HIF-1 inhibition downstream of JVF signalling, whilst not affecting physiological activation of HIF-1 by hypoxia or TPO. This contrasted with Rux, which caused downregulation of HIF-1α levels in both hypoxic and normoxic conditions in JVF cells (Fig.2A[II]). Broader analysis of signalling in JVF mutant UT7/TPO cells demonstrated inhibition of both TPO signalling in JAK2 WT cells and oncogenic JVF signalling, whereas SMI-4a treatment did not deregulate these pathways (Fig.S10). In SET2 and HEL MPN lines (Fig.2B) Rux-treatment significantly downregulated signalling across all pathways analysed, whereas SMI-4a selectively affected levels of HIF-1α. *Ex vivo* expansion of LT-HSCs under SMI-4a demonstrated that PIM1i rescued the JVF het phenotype, returning the LSK/Prog ratios to those of WT LT-HSC cultures (Fig.2C; Fig.S11); this was in contrast to HIF-1 inhibition, which indiscriminately inhibited the LSK fraction regardless of phenotype (Fig.S3A-B). Taken together, these data demonstrate that HIF-1α is stabilised in JVF cells as a result of PIM1 kinase activity, and that inhibition of PIM1 can rescue aberrant HIF-1α activation and JVF HSC phenotypes whilst not altering physiological JAK or HIF-1 activity.

**Figure 2.**
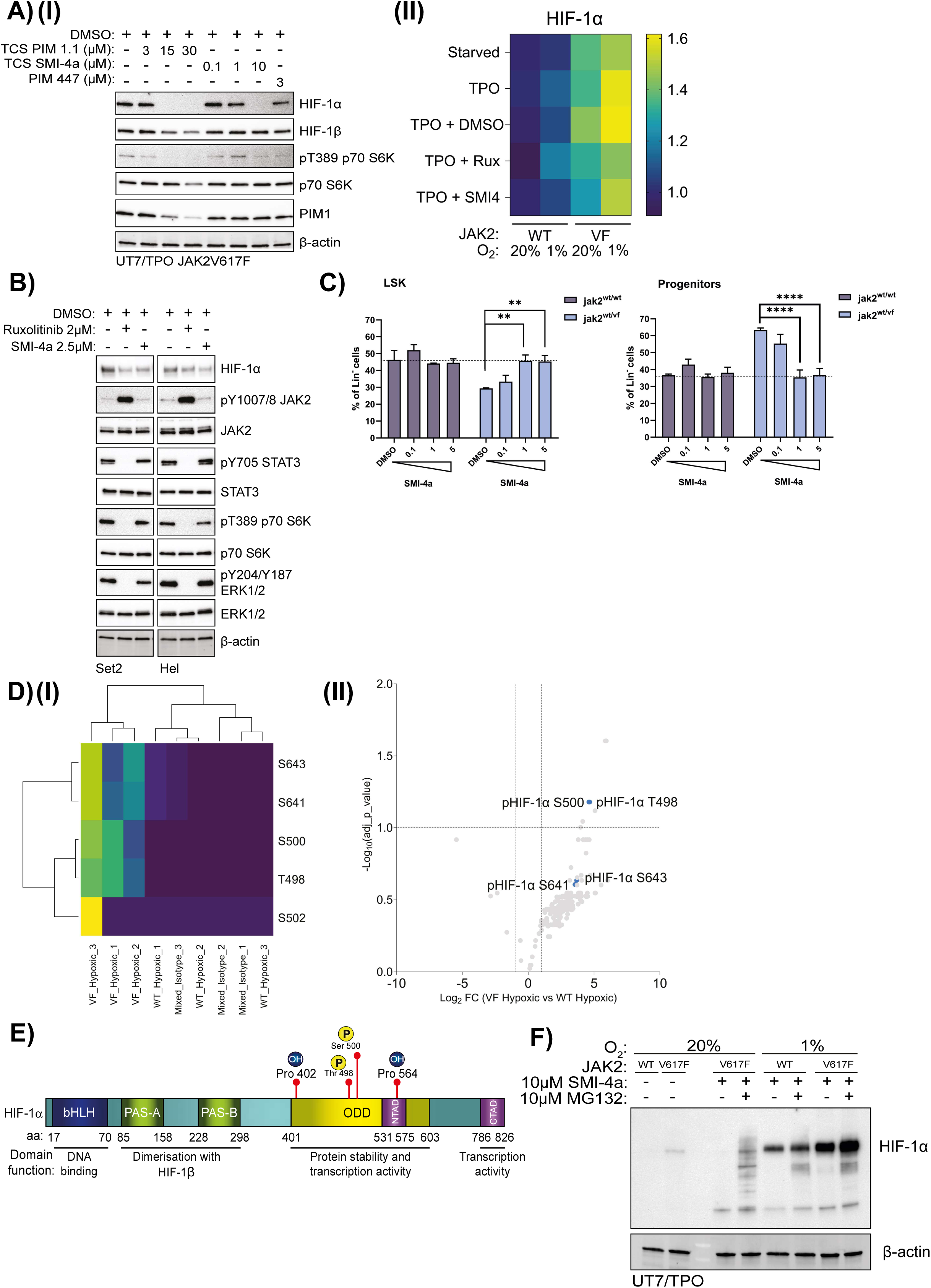
(A)(I) Treatment with a variety of PIM1 inhibitors inhibits JVF-mediated HIF-1α stabilisation. Immunoblot for HIF-1α, HIF-1β, PIM1 and PIM1 substrate (p70 S6K) of JVF UT7/TPO cells starved overnight and treated with the indicated PIM inhibitor for 6hrs at the indicated dose. Immunoblotting carried out using HIF-1α, HIF-1β, pT389 S6, p70 S6K and PIM1 antibodies with β actin as a loading control (n=3). (II) Treatment with SMI-4a restores JVF-stabilised HIF-1α levels closer to those of wildtype TPO-treated cells and leaves wildtype TPO-treated cells unaffected, whilst treatment with Rux ablates both JVF-mediated and wildtype TPO-induced HIF-1α levels. Barcoded phospho-flow cytometry of HIF-1α levels in JVF and wildtype UT7/TPO cells, cultured in normoxic (20% O2) or hypoxic (1% O2) conditions, starved overnight or treated with TPO and SMI-4a, Rux or vehicle control as indicated. Heatmap of mean channel fluorescence intensity (MFI), data normalised based on fold change to mean value of wildtype starved, no TPO, normoxic condition (n=3). (B) Treatment with SMI-4a effectively lowers levels of JVF-stabilised HIF-1α whilst leaving JAK2 signalling cascades untouched, whilst treatment with Rux ablates both JVF-stabilised HIF-1α and JAK2 downstream signalling. Immunoblot for HIF-1α, JAK2, and various kinase cascade constituent proteins downstream of JAK2 of SET2 and HEL cells treated with Rux, SMI-4a or vehicle control as indicated. Performed using HIF-1α, pY1007/8 JAK2, JAK2, pY705 STAT3, STAT3, pT389 p70 S6K, p70 S6K, pY204/Y187 ERK1/2 and ERK1/2 antibodies and using β actin as a loading control (n=3). (C) PIM1 inhibition rescues HSC expansion culture phenotype in JVF cells as determined by proportion of LSK ((lineage negative, Sca-1 positive, c-kit positive) and progenitors in culture, returning the LSK/Prog ratios to those observed in wildtype expansion cultures, whilst leaving wildtype expansion cultures unaffected. Ex vivo expansion of long-term HSCs from wildtype (purple) and JVF heterozygous (blue) mice in the PVA culture system treated with the indicated dose (μM) of SMI-4a or vehicle control and immunophenotyped (by gating strategy following previously described HSC populations [76]) after 14 days of expansion and expressed as percentage of lineage negative cell population (n=3). (D)(I) Two novel JVF-exclusive phosphorylation sites identified on HIF-1α (Thr498 and Ser500), compared to wildtype hypoxia-treated cells. Phosphoproteomic mass spectrometry analysis of hypoxia-treated JVF and wildtype UT7/TPO cell lysate enriched for phosphoproteins by TiO2 and enriched for HIF-1α by immunoprecipitation (n=3). (II) Two novel JVF-exclusive phosphorylation sites identified on HIF-1α (Thr498 and Ser500), compared to wildtype hypoxia-treated cells. Volcano plot of phospho-site signal intensity in hypoxic JVF cell lysate vs hypoxic WT cell lysate plotted vs statistical significance (-log10 adj p value). (E) Novel JVF-exclusive phosphorylation sites (Thr498 and Ser500) fall in HIF-1αs ODD (oxygen-dependent degradation) domain between canonical HIF-1α hydroxylation sites (P402 and P564). Domain diagram of HIF-1α, coloured by domain. Amino acid residues of interest marked in red. (F) Treatment with proteasome-inhibitor MG132 inhibits SMI-4a-mediated destabilisation of JVF-mediated HIF-1α stabilisation. Immunoblot for HIF-1α of JVF and wildtype UT7/TPO cells cultured in normoxic (20% O2) or hypoxic (1% O2) conditions treated with SMI-4a, MG132 or a combination of the two as indicated, using HIF-1α antibody and using β actin as a loading control (n=2).

We next performed phospho-proteomic mass spectrometry analysis of UT7/TPO WT/JVF cells exposed to hypoxia; HIF-1α phosphorylation was detected at the canonical hypoxia/ERK sites (Ser641/643) in WT cells, whereas in hypoxic JVF cells an additional phosphorylation couplet was detected at Thr498/Ser500 (Fig.2D(I)-[II]). The Ser500 site is species-conserved, demonstrating its significance as a phospho-regulatory residue (Fig.S12). Interestingly, no phosphorylation at the Thr455 site was detected in any condition, suggesting that the mechanism of PIM1-mediated HIF-1α stabilisation downstream of JVF is distinct from that observed in prostate cancer cells^38^. The location of Thr498/Ser500 within the ODD and proximity P402/564 hydroxylation sites (Fig.2E; Fig.S13) suggested that the mechanism of PIM1-mediated HIF-1α stabilisation may be similar; we therefore treated WT/JVF UT7/TPO cells with the proteasome inhibitor MG-132 in combination with SMI-4a (Fig.2F). Combined SMI-4a/MG132 treatment re-stabilised HIF-1α protein, demonstrating that PIM1-mediated stabilisation of HIF-1α downstream of JVF signalling occurs via inhibition of canonical normoxic proteasomal degradation.

### HIF-1α assembles a non-canonical transcription complex in JAK2V617F cells

We next sought to determine whether the non-canonical phosphorylation-mediated stabilisation of HIF-1 in JVF cells altered its transcription activity. HIF-1α phosphorylation is known to alter its interaction with co-regulatory proteins^46,47^, we therefore wanted not only to capture the target genes of HIF-1 in JVF cells, but also the transcription co-regulators. We therefore developed a dual-output chromatin-interactome assay (**DOCIA**), integrating the principles of ChIPseq and qPLEX-RIME (quantitative multiplexed Rapid Immunoprecipitation Mass spectrometry of Endogenous proteins), to quantitatively analyse both the genomic DNA and protein interactors of HIF-1α from a single input sample (Fig.3A[I]). HIF-1α ChIP was performed on WT/JVF BaF3/hMPL normoxia (20%)/hypoxia (1%, 6h) cells. First, we observed that despite equal input chromatin loading and HIF-1α protein between WT/JVF conditions (Fig.S14) on to the ChIPs, we observed significantly higher HIF-1α protein levels (MS) at the chromatin in JVF cells compared to WT cells (Fig.3A[II]). Unexpectedly, this was coupled with a significant reduction in DNA associated with HIF-1α in JVF cells (NGS) compared to WT, particularly evident in the VF_Nx cells.

**Figure 3.**
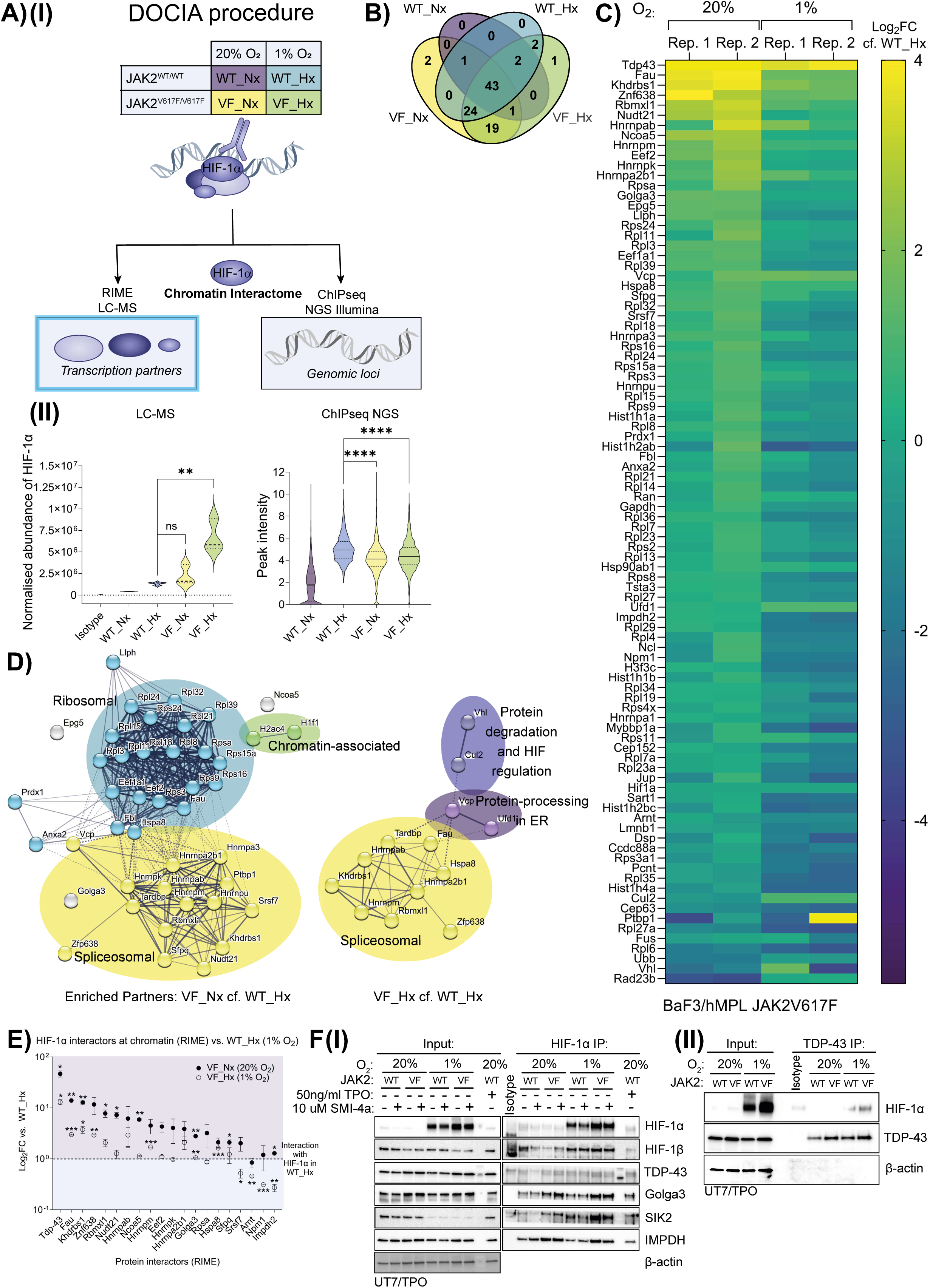
(A)(I) Diagram of DOCIA method: wildtype or JVF hMPL BaF3 cells cultured in normoxic (20% O2), or hypoxic conditions (1% O2) have their nuclear material crosslinked to retain DNA-bound proteins, fragmented and HIF-1α containing fragments immunoprecipitated by the addition of HIF-1α antibody. The sample is then divided for concurrent analysis by both ChIP-seq (to determine HIF-1αs genomic binding loci) and mass spectrometry (RIME-Rapid immunoprecipitation mass spectrometry of endogenous protein) (to determine HIF-1αs associated transcription cofactors). (II) JVF causes significantly increased abundance of HIF-1α protein at the chromatin in hypoxic conditions yet is significantly less bound to it compared to wildtype cells cultured in hypoxia in both normoxia and hypoxia. RIME-mass spec analysis of HIF-1α abundance at the chromatin (left) and ChIPseq analysis of the general peak intensity of chromatin bound by HIF-1α (right) in wildtype and JVF hMPL BaF3 cells cultured in either normoxic or hypoxic conditions. Coloured by condition as described in Fig.4A(I) (n=3). (B) Whilst several cofactors are found to be common between all conditions, a number are identified as unique to JVF in both normoxia and hypoxia. Venn diagram displaying number of cofactors unique to each condition shown as determined by RIME-mass spec. Coloured by condition as described in Fig.4A(I) (n=3). (C) RIME-mass spec-identified HIF-1α cofactors displayed as a heatmap by binding intensity (Log2FC), normalised to HIF-1α protein levels. Wildtype and JVF hMPL BaF3 cells cultured in either normoxic (20% O2) or hypoxic (1% O2) conditions and subjected to RIME-mass spectrometry wing of DOCIA procedure (n=3). (D) Distinct JVF-unique functional protein clusters are enriched in HIF-1α cofactor analysis. STRING analysis was performed on RIME-mass spec protein hits. For each treatment condition, structural and non-mouse proteins were discarded, RIME hit proteins were sorted by geomean (relative to HIF-1α) and those less than 0.01 discarded. The remaining proteins were then run through the STRING analysis online database and clustered via MCL clustering (stringency 3). Resulting connections plotted by confidence (cutoff ‘medium’ – 0.04). (E) Key cofactors were identified as uniquely up or downregulated in association with HIF-1α in JVF cells in both normoxia and hypoxia. Wildtype and JVF hMPL BaF3 cells cultured in either normoxic (20% O2) or hypoxic (1% O2) conditions and subjected to RIME-mass spectrometry wing of DOCIA procedure, the top 20 most significantly up or down regulated proteins in both JVF normoxic and hypoxic cells plotted on chart by Log2FC relative to wildtype (n=3). (F)(I) Key cofactors identified by DOCIA validated by HIF-1α immunoprecipitation. Immunoblot of HIF-1α, HIF-1β and key JVF-unique cofactors identified by RIME mass spec in HIF-1α-immunoprecipitated wildtype and JVF UT7/TPO cells cultured in either normoxic (20% O2) or hypoxic (1% O2) conditions, treated with or without SMI-4a as shown and with TPO-treated wildtype normoxic cells as additional control. Input on left, IP on right. Antibodies for HIF-1α, HIF-1β, TDP-43, Golga3, SIK2 and IMPDH used with β-actin used as loading control (n=2). (II) The JVF-unique cofactor TDP-43, identified by DOCIA, validated by TDP-43 immunoprecipitation of HIF-1α. Immunoblot of HIF-1α and TDP-43 in TDP-43-immunoprecipitated wildtype and JVF UT7/TPO cells cultured in either normoxic (20% O2) or hypoxic (1% O2) conditions, with HIF-1α and TDP-43 antibodies used and with β-actin as loading control. Input on left, IP on right (n=2).

Analysis of the protein interactors of HIF-1α at the chromatin in WT_Hx cells identified known co-regulator interactors including HIF-1β/ARNT, Hsp90 and Ncoa5 (Supplementary Tables S1-S2)^47^. Common cofactors associating with HIF-1α across all conditions included HIF-1β/ARNT, Ncoa5, Epg5, polyubiquitin-B, centrosomal protein Cep63 and Gapdh. 94 proteins were found to be significantly enriched with HIF-1α relative to isotype control across the conditions, of which 43 were common to all four conditions and 24 to those conditions with stabilised/active HIF-1a (WT_Hx, VF_Nx, VF_Hx) (Fig.3B and Supplementary Table S2). Strikingly, we detected 19 binding partners that are uniquely associated with HIF-1α in cells harbouring the JVF mutation, whereas we did not detect any wildtype-specific factors. Exposure of a given genotype to hypoxia also did not substantially alter the associated factors, with only 3 proteins detected only in hypoxic mutant cells (VF_Hx/+WT_Hx): these included two ribosomal components (Fbl, Rps3a) and the DNA damage repair protein Rad23.

Quantitative analysis of differences in HIF-1α cofactors between JVF/WT cells, whereby samples were first normalised against HIF-1α abundance (Fig.3A[II], Supplementary Table S1) and then against WT_Hx interaction (Fig.3C), illuminated key patterns of JVF-unique protein association, such as an increased association of HIF-1α with cofactor TDP-43 and decreased association with protein markers of degradation (ubiquitin; VHL) lending evidence to our earlier mechanistic findings implicating inhibition of HIF-1α proteasome-degradation in JVF cells. In VF_Hx conditions, the proteactome of HIF-1α was substantially reduced, despite greater levels of HIF-1α relative to WT_Hx and VF_Nx (5.42 and 3.15-fold greater respectively), suggesting a significant alteration in HIF-1α function in hypoxic JVF cells. Most interestingly, an emergent group of spliceosomal factors were uniquely enriched in interaction with HIF-1α in JVF cells under both conditions, (including TDP-43, KHDRBS1/Sam68), along with ribosomal factors in normoxic treated cells (Fig.3D; Fig.S15A-B). JVF-unique interacting partners identified through DOCIA were then validated by endogenous co-immunoprecipitation assays (Fig.3E; Fig.3F(I)-[II]). This confirmed the interaction of TDP-43, GOLGA3, SIK2 and IMPDH with endogenous HIF-1α. Phosphoproteomic investigation of HIF-1α phosphorylation (Fig.2D[I]), also identified GOLGA3 as a phosphorylated co-immunoprecipitated protein, with several distinct phospho-sites detected in JVF cells (Fig.S16). Together, these data identify that differential HIF-1 transcription complexes form as a direct result of the JVF mutation, comprising non-traditional co-regulatory proteins and significant enrichment of pre-mRNA processing factors.

### HIF-1 is genomically redistributed in JAK2V617F cells

Continuing our DOCIA methodology (Fig.4A), we next performed NGS upon genomic DNA immunoprecipitated with the HIF-1LJ complexes analysed in Fig.3. First, we identified a canonical HIF-1 signature^48,49^ in WT_Hx conditions (Fig.S17), including classical target genes *Vegfa*, *Slc2a1, Eno1* and *Gapdh* (Fig.S18). Globally, we observed a reduction in median peak intensity (MPI) associated with HIF-1α in JVF cells (Fig.4B, Fig.S19), comprising a population of loci of low intensity (MPI<2), unobserved in WT_Hx condition (despite higher HIF-1α protein levels (Fig.3A[II]). Concomitant with this, we observed very few HIF-1α loci that were unique to JVF mutant cells, rather the binding loci were a subset of those bound by HIF-1α in WT_Hx conditions (Fig.S20). These data indicate that the DNA-binding capacity and/or the proximity of HIF-1α to DNA is reduced in JVF mutant cells at particular sites compared to hypoxic WT JAK2 cells.

**Figure 4.**
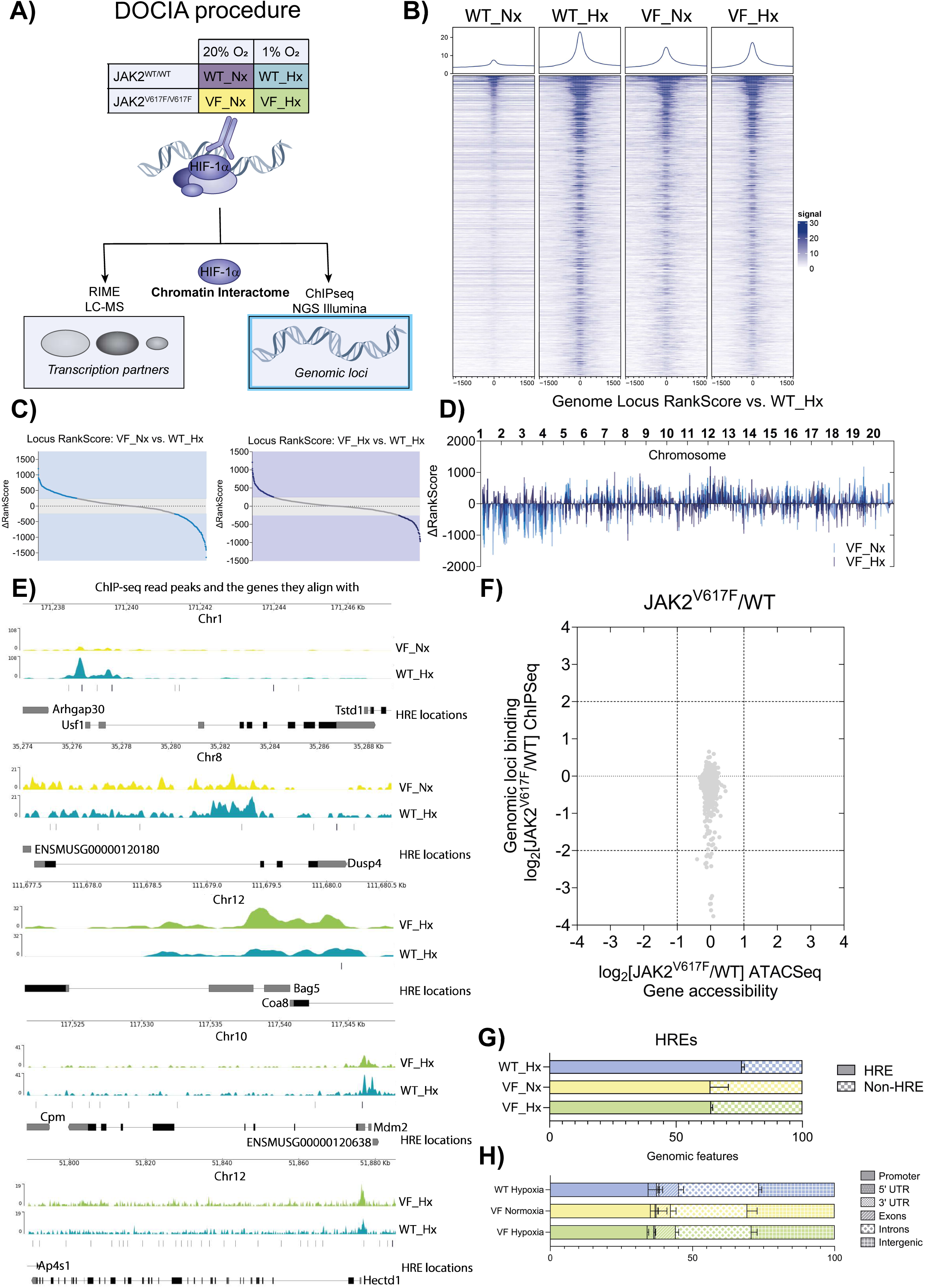
(A) Diagram of DOCIA method: wildtype or JVF hMPL BaF3 cells cultured in normoxic (20% O2), or hypoxic conditions (1% O2) have their nuclear material crosslinked to retain DNA-bound proteins, fragmented and HIF-1α containing fragments immunoprecipitated by the addition of HIF-1α antibody. The sample is then divided for concurrent analysis by both ChIP-seq (to determine the genomic binding loci of HIF-1α) and mass spectrometry (RIME-Rapid immunoprecipitation mass spectrometry of endogenous protein) (to determine the associated transcription factors of HIF-1α). (B) Weaker overall binding of HIF-1α to chromatin observed in normoxic and hypoxic JVF cells compared to hypoxic-treated wildtype cells. Heatmap of normalised ChIPseq signal for the entire genome from wildtype and JVF hMPL BaF3 cells cultured in normoxic (20% O2) or hypoxic (1% O2) conditions and subjected to the ChIP-seq wing of the DOCIA procedure, as displayed by median peak intensity (n=3). Each row of the heatmap is a genomic region, centred to peaks of binding signals. Binding is summarized with a gradient colour code key with 0 representative of no binding (white) to maximum binding 30 (blue). Plots at the top of the heatmaps show the median signal at genomic regions centred at peaks of binding signals. (C) Whilst the genomic loci bound by HIF-1α in JVF cells are minimally altered when compared to wildtype hypoxic-treated cells, the priority in which those sites are bound by HIF-1α is significantly altered in both normoxic and hypoxic JVF cells, with a number of loci whose HIF-1α binding priority relative to WT_Hx binding is altered by more than a quartile in both VF_Nx and VF_Hx conditions. Δrank-score analysis of HIF-1α-bound genomic loci for VF_Nx (left) VF_Hx (right) denoting loci higher or lower ranked in HIF-1α binding priority than WT_Hx, as determined by ChIP-seq, plotted in order of score, with those loci with Δrank-score greater than a quartile highlighted. (D) Across the mouse genome, the priority of HIF-1α binding appears lessened between chromosomes 1-4 and sites at chromosomes 8 and 14 appear prioritised in VF_Nx when compared to WT_Hx whilst the priority of HIF-1α binding appears enhanced in the region of chromosome 12 when compared to WT_Hx. Δrank-score analysis of HIF-1α-bound genomic loci for VF_Nx (light blue) VF_Hx (dark blue) denoting loci higher or lower ranked in HIF-1α binding priority than WT_Hx, as determined by ChIP-seq, plotted by genomic location (chromosome). (E) Key genes found to be differently bound by HIF-1α in VF_Nx and VF_Hx compared to WT_Hx include a loss in Usf1 and a gain in Dusp4 HIF-1α binding in VF_Nx and a loss in Mdm2 and a gain in Bag8, Coa8 and Hectd1 HIF-1α binding in VF_Hx. Gene tracks of genomic loci where significant loss or gain of HIF-1α binding as determined by ChIP-seq can be observed in VF_Hx (green) or VF_Nx (yellow) compared to WT_Hx (blue). (F) Cross analysis of ChIP-seq data with scATAC-Seq data demonstrates no correlation between gene loci accessibility and HIF-1α binding in JVF compared to wildtype. Plot of genome-wide gene accessibility (as derived from [52] Supplementary Figure 4 ‘Differential gene accessibility score in HSC and HSCMY clusters in untreated patients with myelofibrosis.’) in primary JVF-mutation carrying patient cells plotted against genome-wide genomic loci binding (as determined by ChIP-seq of hMPL BaF3 VF_Hx cells). (G) Analysis of ChIP-seq-derived HIF-1α binding loci shows reduced binding to canonical HREs (Hypoxic Response Elements) in JVF cells in both normoxia and hypoxia. HIF-1α HRE binding analysis of wildtype and JVF hMPL BaF3 cells cultured in normoxic (20% O2) or hypoxic (1% O2) conditions and subjected to the ChIP-seq wing of the DOCIA procedure, shown as percentage of HIF-1α binding loci localised to HREs. (H) Analysis of ChIP-seq-derived HIF-1α binding loci shows increased binding in intergenic regions of the DNA in JVF cells in both normoxia and hypoxia and decreased exon binding in normoxia. Genomic feature binding analysis of wildtype and JVF hMPL BaF3 cells cultured in normoxic (20% O2) or hypoxic (1% O2) conditions and subjected to the ChIP-seq wing of the DOCIA procedure, shown as percentage of HIF-1α binding loci localised to the indicated genomic features.

We next calculated the priority distribution of HIF-1 across the genome within a given cellular context, applying Δrank-score for VF_Nx/VF_Hx relative to WT_Hx. First, we observe global priority redistribution of HIF-1 in JVF cells compared to WT_Hx, with Δrank-score >Q1 of 42%/26% of sites in VF_Nx/VF_Hx, respectively (Fig.4C). Analysing the chromosomal distribution of intensity ranking, a population of depreciated HIF-1 bound sites in VF_Nx were evident in chromosomes 1-4 and prioritised sites at chromosome 8 and 14 (Fig.4D; Fig.S21); notable genetic loci losses and gains included *Usf1* and *Dusp4*, respectively (Fig.4E). In VF_Hx cells HIF-1α binding was overall enhanced on chromosome 12 (Fig.4D; Fig.S21); these sites included *Bag5*, *Coa8, Mdm2* and *Hectd1* (Fig.4E). To determine whether chromatin accessibility alterations mediated by JVF signalling was a factor in the redistribution in HIF-1α, we cross-analysed VF_Nx/VF_Hx ChIPseq data with a JVF scATAC-Seq dataset (Fig.4F; Fig.S22)^50^. No significant difference in accessibility at HIF-1α binding loci was observed, suggesting that HIF-1α redistribution was not mediated by site accessibility. Feature analysis of HIF-1 bound loci demonstrated a 14% reduction in hypoxia-responsive elements in VF_Nx/VF_Hx compared to WT_Hx (Fig.4G) and a modest increase in binding in intergenic regions and introns in VF_Nx cells (Fig.4H).

### JAK2VF-HIF-1 gene-signatures diverge from canonical HIF-1 targets and correlate with MPN disease progression and survival

Next, gene-signatures were generated for enriched sites in WT_Hx (relative to WT_Nx), VF_Nx and VF_Hx (relative to WT_Hx) (Fig.5A). Gene ontology (GO) pathway analysis of WT_Hx identified canonical HIF-1 hallmark gene pathways^48,49^, including glycolysis and response to oxidative stress and hypoxia. (Fig.5B[I]). Non-canonical pathways were evident in the VF_Nx signature, including DNA metabolism, DNA damage response (DDR), and RNA processing, and reduction of glycolysis pathways (Figure 5B[II]); strikingly, the VF_Hx signature GO was even further separated from WT_Hx pathways, with cell cycle progression and checkpoint pathway genes demonstrating the greatest contribution to the signature (Fig.5B[III]). Furthermore, VF_Nx/VF_Hx signatures contained only 9/2 genes, respectively, of the HIF gene metascore (core 48 genes highly conserved across cell/cancer types exposed to hypoxia^3^) (Fig.S23), underscoring the functional divergence incurred by the modality of HIF-1 activation in JVF mutant cells.

**Figure 5.**
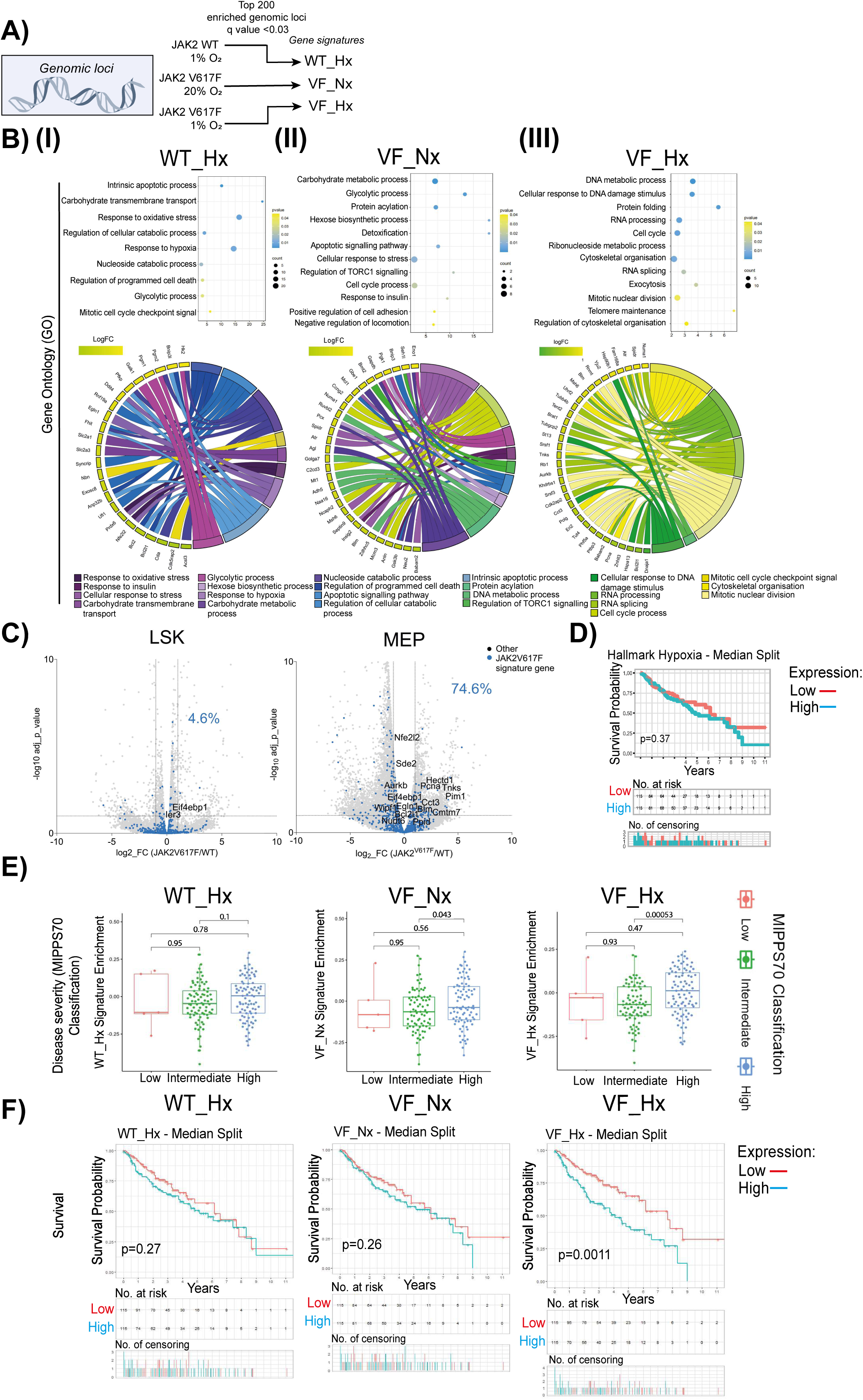
(A) Diagram of derivation of ‘gene signatures’: Top 200 enriched genomic loci from HIF-1α ChIP-seq analysis with q<0.03 for each condition selected for further analysis. B)(I) GO (gene ontology) analysis of WT_Hx ‘gene signature’ shows key canonical HIF-1 pathway associations. GO data of WT_Hx ‘gene signature’ displayed both as enrichment bubble plot (top) and chord plot (bottom). Enrichment bubble plot plotted in descending order of statistical significance as designated by p-value (from 0.01 to 0.04) with ‘bubble’ size indicating number of genes associated with the indicated function. Enrichment chord plots plotted by gene within signature in order of fold change (LogFC) and its associated GO function (colour-coded). (II) GO (gene ontology) analysis of VF_Nx ‘gene signature’ shows loss of key canonical HIF-1 pathway associations (such as glycolysis) and gain of novel pathway associations never before ascribed to HIF-1α including DNA metabolism, DNA damage response and RNA processing. GO data of VF_Nx ‘gene signature’ displayed both as enrichment bubble plot (top) and chord plot (bottom). (III) GO (gene ontology) analysis of VF_Hx ‘gene signature’ shows more extensive loss of key canonical HIF-1 pathway associations and gain of novel pathway associations never before ascribed to HIF-1α including RNA splicing, RNA processing, cell cycle progression and checkpoint pathway genes. GO data of VF_Nx ‘gene signature’ displayed both as enrichment bubble plot (top) and chord plot (bottom). (C) The transcripts of 74.6% of the genes in the VF_Hx gene signature are detectable in mouse MEP (megakaryocytic-erythroid progenitor) cells in bulk RNAseq experiments, whilst transcripts of only 4.6% of the signature are detectable in mouse LSK (lineage negative, Sca-1 positive, c-kit positive) cells. Volcano plots of bulk RNAseq from mouse primary haematopoietic stem and progenitor cells (HSPCs) from wildtype (n=3) and JVF homozygous (n=3) C57/BL6 mice, (LSKs top and MEPs bottom). VF_Hx gene signature transcripts highlighted in blue with transcripts with significance higher than −log10 adj p value of 1 labelled. (D) ‘Hallmark hypoxia gene set’ shows no correlation with JVF MPN patient cohort survival. (E) Neither WT_Hx nor VF_Nx ‘gene signatures’ show correlation with MPN patient disease severity as defined by MIPSS70 classification whereas the VF_Hx signature does. MPN patient cohort (172 JVF mutation carrying MPN patients) expression enrichment of WT_Hx, VF_Nx and VF_Hx gene signatures plotted by MIPSS70 (Mutation-Enhanced International Prognostic Score System for transplantation eligible-aged patients with overt PMF) classification. The score takes into account haemoglobin <10 g/dL (1 point), leukocyte count >25×109/L (2 points), platelet count <100×109/L (2 points), circulating blasts ≥2% (1 point), MF-2 or higher BM fibrosis grades (1 point), presence of constitutional symptoms (1 point), absence of CALR type-1 mutation (1 point), and presence of HMR mutations (ASXL1, EZH2, SRSF2, and IDH1/2; 1 point for single mutation and 2 points for ≥2 HMR mutated genes) as independent prognostic factors of inferior OS. The MIPSS70 classified the MF patients into three groups: low-risk (0–1 point), intermediate-risk (2–4 points), and high-risk (≥5 points). (F) MPN patient cohort expression enrichment of neither the WT_Hx nor the VF_Nx ‘gene signatures’ show correlation with JVF MPN patient cohort survival whereas patient enrichment of VF_Hx ‘gene signature’ expression show significant correlation with poorer survival outcomes. MPN patient cohort survival curves subdivided into populations with high (blue) or low (red) expression of WT_Hx, VF_Nx or VF_Hx gene signatures with significance of difference between the populations shown on graph (p-value). Remaining patients in each population shown in table below with number of censoring (i.e. number of patients leaving the study whose fate is unknown).

To investigate whether the redistribution of HIF-1 in response to JVF signalling was reflected in the transcriptome, we next interrogated the expression of the VF_Nx / VF_Hx signatures in JVF HSPCs RNAseq (Fig.5C)^39^. This revealed 74.6% of VF_Hx signature genes were differentially expressed in JVF MEP cells, compared to only 4.6% in LSK cells. Interestingly, we observed that a significant proportion of these genes were downregulated in MEPs. Expression was not mediated by genotype-induced chromatin accessibility differences; cross analysis with ATACseq datasets showed no correlation with VF_Hx signature gene expression (Fig.S24; Fig.S25).

We next interrogated the HIF signatures in a 172-patient cohort of JVF-positive MPNs with matched RNAseq and disease progression/survival data^51^, to determine whether adverse HIF-1 transcriptional output in response to JVF mutation conferred pathophysiological effects in MPN disease. First, we discovered no correlation between the WT_Hx signature or a classical hypoxia signature (Hallmark_Hypoxia) in either disease progression or survival of these patients (Fig.5D; Fig.5E-F). However, interrogation of the VF_Nx/ VF_Hx signatures demonstrated a significant correlation between expression and disease progression from intermediate to high MIPSS70 score (Fig.5E)^52,53^. Correlation of the VF_Hx signature with disease severity was reflected in the overall survival of the cohort, for whom those in the upper median group of VF_Hx gene signature expression (VF_Hx ^HIGH^) demonstrated significantly worse overall survival, with a median survival of 4 years compared to 8 years for VF_Hx^LOW^ patients (Fig.5F).

### JAK2V617F-activated HIF-1 target genes contribute to disease progression and outcome

To identify whether particular genes in the VF_Hx signature were responsible for survival outcome, we next analysed variable expression of these genes across a range of biologically motivated genesets, including ATAC-seq^50^, bulk murine RNAseq^39^, scRNA-seq from JVF MF patients^40^ and our 172-JVF MPN patient cohort (Fig.6A). First, we observed that the JVF-HIF-1 target genes significantly associated with disease severity (MIPSS70 score) and overall survival of JVF patients, were differentially expressed within the disease-driving megakaryocyte progenitor populations in both humans (MkPs) and mice (MEPs) (Fig.6B(I)-[II]). A key emergent cellular pathway was the DNA damage response^54^, with the JVF-HIF-1 target genes *PCNA, BLM, AURKB, EIF4EBP1 and BCL2XL* (Fig.S26) upregulated in MEPs/MkP and associated with disease progression/survival. JVF-HIF-1 genes that were globally upregulated throughout HSPC clusters, for example *TNKS, HECTD1, WIPF1, PPID* (Fig.S27) were not correlated with disease progression or survival. Additionally, we noted a small cohort of genes that were significantly upregulated in more primitive progenitor populations in patients (HSPC3), that were inversely correlated with disease severity and survival, including *IER3*, which regulates cell cycle/apoptosis decisions (Fig.S28)^55^. These data support previous findings that the disease-specific expanded MkP population drives disease in JVF MF patients^50^, and also identify that disease co-opted HIF-1 downstream of JVF signalling induces DNA damage response genes that contribute to MPN disease progression and patient survival.

**Figure 6.**
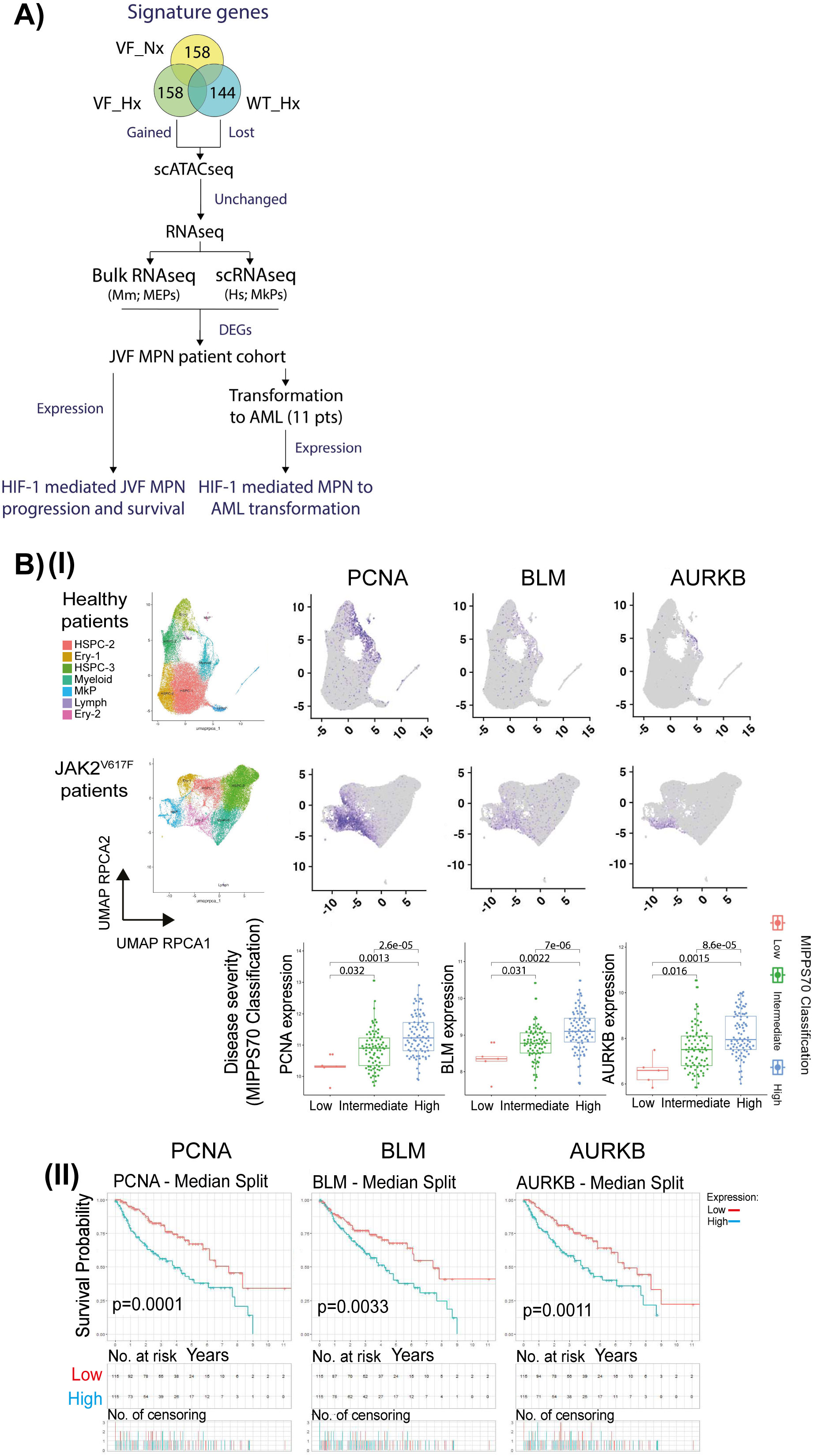
(A) Diagram illustrating identification of specific genes responsible for differential survival in patients with high VF_Hx signature expression compared to patients with high VF_Nx signature expression. Genes unique to VF_Hx signature were run through JVF MPN patient scATACseq, bulk mouse RNAseq and scRNAseq human datasets, any genes significantly upregulated in all three were run through the MPN patient cohort dataset and remaining genes associated with poorer patient survival examined. (B)(I) Genes identified as responsible for differential survival in MPN patients with high VF_Hx expression include genes associated with DNA damage response and are found to be upregulated specifically in patient MkP cell populations. High expression of these individual genes is also correlated with MPN patient disease severity. UMAP scRNAseq plots of control (top row) and myelofibrosis (bottom row) HSPCs with transcripts of genes of interest (PCNA, BLM and AURKB) highlighted in purple. The correlation of the individual genes of interest with MIPSS70 MPN patient classification displayed below. (II) High expression of the individual genes PCNA, BLM and AURKB is associated with significantly poorer MPN patient survival outcomes.

### JAK2VF-HIF-1 regulon correlates with transformation of MPNs to AML

JVF positive MPN patients who develop myelofibrosis are at risk of spontaneous conversion to acute myeloid leukaemia; however, the molecular mechanisms underpinning this pathobiological event are not well characterised^7^. We therefore applied the above sequential analysis to 11 patients from the JVF MPN cohort that transformed to AML, analysing expression of JVF-HIF-1 variable genes in matched pre-/post-transformation samples. As expected, no significant difference in expression of the gene signatures as a whole was observed in these patients pre-/post-transformation (Fig.S29), we identified a subset of HIF-1 genes (13) that were significantly associated with transformation (Fig.7A). DNA damage response genes were pre-eminent as upregulated within this subset, whereas downregulated pathways included negative feedback of the hypoxic response (*EGLN1*) and regulators of glucose metabolism (*IER3, HK2*). Analysis of these genes as a unified regulon demonstrated an extremely strong correlation with disease progression (MIPSS70 classification), myelofibrosis (overt, post-ET and post-PV), and significantly worsened survival (Fig 7B). Together, these results uncover a hitherto unidentified, non-canonical HIF-1 regulon in JVF-mutant MPNs that drive progression, leukemic transformation and survival outcome for patients.

**Figure 7.**
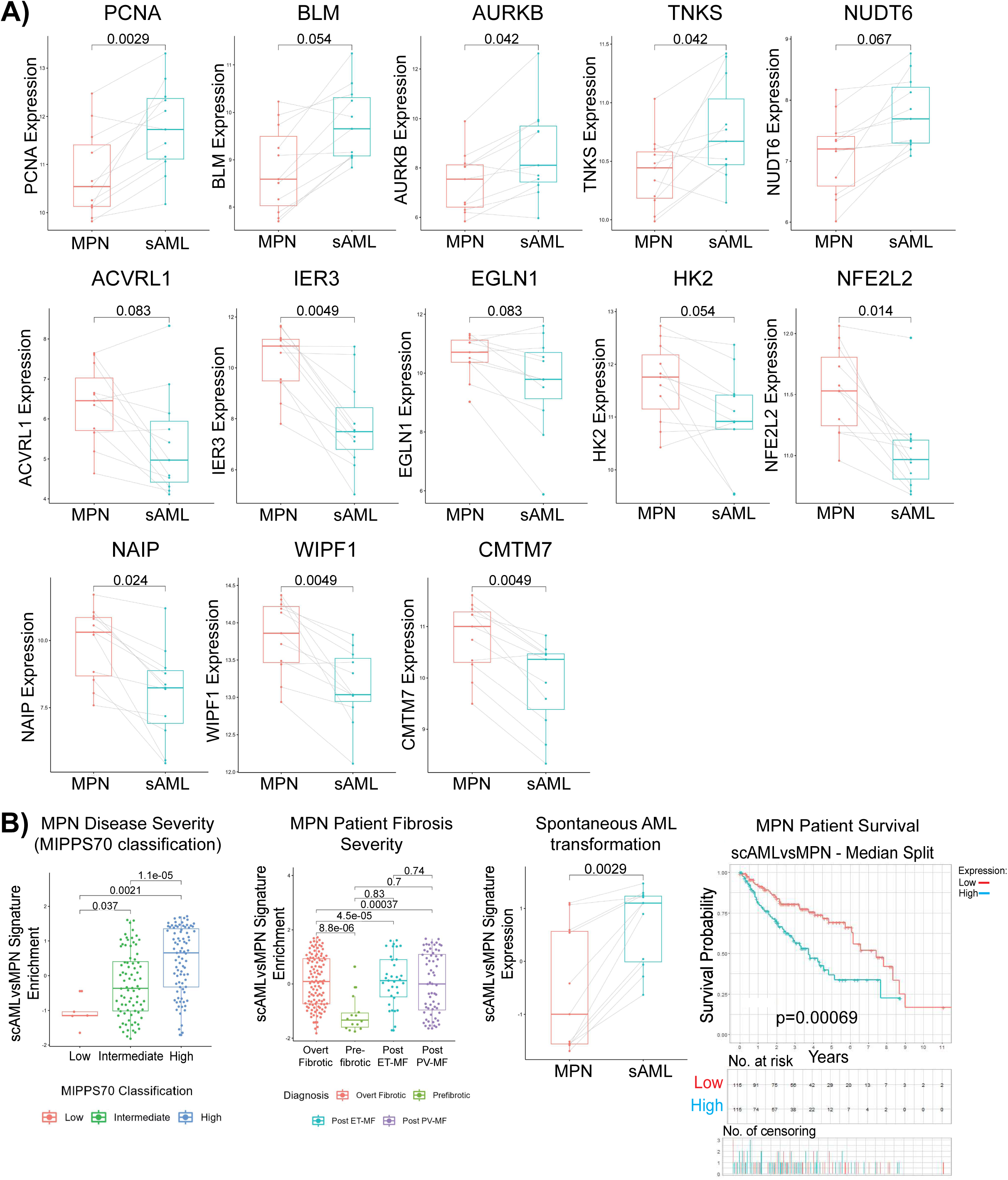
(A) Of the genes identified by the process outlined in Fig.7A, the high expression of a subset of them were found to be associated with increased risk of spontaneous AML transformation. Paired transcript expression of the genes of interest pre (red) and post (blue) spontaneous AML (sAML) transformation plotted by MPN patient. The top and middle rows of the figure illustrate genes whose expression were elevated in patients post-spontaneous AML transformation and the bottom row illustrates genes whose expression levels were lowered upon spontaneous AML transformation. (B) Taken together as a novel ‘sub-signature’, high expression of the genes that show elevated expression in MPN patients who suffered spontaneous AML transformation are significantly associated with MPN patient disease severity, increased myelofibrosis, AML transformation and poorer patient survival. From left to right, MPN patient cohort expression enrichment of the spontaneous AML transformation associated ‘gene signature’ plotted by MIPSS70 classification, expression of the signature plotted by severity of patient myelofibrosis, expression of the signature in patients pre and post spontaneous AML transformation and survival of patients stratified into those with high or low expression of the signature.

## Discussion

Through direct experimental analysis of pathologically activated HIF-1 transcription complexes, our study reveals that JAK2V617F signalling reprograms HIF-1 function, activating a non-canonical regulon, driving MPN disease. Our study resolves long-standing questions regarding the multifunctionality of HIFs within neoplasia^4,5,9,56^ through the identification that the HIF-1 regulon is substantially altered by the modality of HIF-1 activation.

Our findings prove PIM1 is responsible for oxygen-independent activation of HIF-1 in JVF MPNs via phosphorylation of the ODD and inhibition of proteasomal degradation via PHD/VHL. PIM1 was previously identified as conferring phenotypic effects in murine and *in vitro* models of JVF malignancy^30,57,58^ and elevated PIM1 levels are reported as disease driving in AML^59,60^, mantle cell lymphoma^61^ and diffuse large B-cell lymphoma^62,63^. Further characterisation of the effects of Thr498/Ser500 phosphorylation on HIF-1α structure/function, and investigation of PIM1-mediated activation of HIF-1 in other haematological/solid cancers are warranted. Whereas HIF-1 inhibitors have been unsuccessful as cancer therapeutics^64,65^, our study reveals PIMi eradicates HIF-1α in JVF cells whilst not impairing TPO-/hypoxia-mediated HIF-1 activation, and corrects aberrant lineage committal decisions within JVF HSPCs; these data define downstream mechanistic effects of PIM1i, and provide proof-of-principle that therapeutic targeting of pathological HIF-1 via context-specific activatory mechanisms is achievable.

Our findings heavily implicate HIF-1 as performing transcript-regulatory role beyond transcription initiation in JVF cells: HIF-1a demonstrated reduced total association with DNA, enhanced binding in intron/exon regions, enhanced interaction with post-transcriptional RNA processing factors, and target-gene expression that was both up- and down-regulated. A subset of transcription factors has been described to regulate pre-mRNA processing via imprinting and regulation of elongation rates^66–69^. In breast cancer cells, HIF-1α binds to TRIM28/DNAPK to release paused RNA Pol II, regulating the rate of elongation^46^. Our findings further implicate HIF-1 as regulating pre-mRNA processing in JVF-HSPCs to produce the observed disease-driving regulon of suppressed and activated genes in MPN disease, however further investigation is required to fully elucidate this function.

Within the JVF-HIF-1 regulon, a group of DNA damage response (DDR) genes were significantly associated with patient outcome. JVF-HSPCs are known to accumulate spontaneous DNA damage^70–73^ and activate the DDR ^70,72^. Our findings identify that malignant co-option of HIF-1 facilitates this in part via upregulation of nucleotide excision repair^74^, homologous recombination^75,76^, and mitotic checkpoint genes^77^; each of these are also significantly associated with AML transformation. The DNA damage response was also pre-eminent in a subgroup (13 genes) of the JVF-HIF-1 regulon that is significantly associated with spontaneous leukaemic transformation of JVF MPN patients. To our knowledge, these JVF-HIF-1 DDR genes have not previously been associated with MPN leukaemic transformation and warrant further investigation.

Our findings identify a substantial discrepancy between the canonical HIF-1 regulon and those induced by oncogenic signalling pathways in the context of JVF MPNs, providing direct evidence that functional delineation of the essential homeostatic roles of HIF-1 within normal physiology from those roles co-opted by malignancy is possible. Our study rationalises future investigations into the malignant co-option of other TFs in JVF MPNs, and of HIF-1 in the pathogenesis of other haematological and solid cancers.

## Author Contributions

D.K. and R.E. designed experiments, performed research, analysed results and wrote the manuscript; S.B., A.G.X.Z., J.J.F.M, R.T.G., G.W., N-M.B, K.A.W., C.A.H., J.L. H.M.K., G.C., B.D., J.C., B.L.F., A.K.F., D.G.K., B.P., A.C.W., A.N.H., I.S.H., A.S.M., J.E.D. participated in the research; V.G. contributed the clinical data; K.S.B. conceived the project, designed and performed experiments, supervised the research and wrote the manuscript; all authors reviewed and approved the manuscript.

