## Supplemental Information for "HIF-1 activated by PIM1 assembles a pathological transcription complex and regulon that drives JAK2V617F MPN disease"

**Supplementary Information**

**Supplementary Methods and Method Figures:**

**Dual Output Chromatin Interactome Assay (DOCIA) extended method**

$5x10^7$ BaF3 hMPL hJAK2 WT or BaF3 hMPL hJAK2 VF cells were incubated at the indicated $O_2$ concentrations for 6 hours. Cells were fixed by addition of 2mM Disuccinimidyl glutarate (DSG) (Chem Cruz - Santa Cruz) for 20 minutes with agitation, pelleted by centrifugation at 500xg for 5 minutes RT, and then resuspended into 1% formaldehyde (Cell Signalling Technology) for 10 minutes at room temperature (RT), then neutralised by the addition of 10x Glycine for 5 minutes at RT. Sample processing and immunoprecipitations were performed as described previously [41,42]. **Chromatin-Immunoprecipitation:** Samples were then processed using SimpleChIP® Enzymatic Chromatin IP Kit, Magnetic Beads (Cell Signaling Technology) according to the manufacturer's protocol. ChIP was performed with HIF-1α (D1S7W) XP® Rabbit mAb (Cell Signaling Technology). DNA was isolated using a ChIP DNA Clean & Concentrator (Zymo Research). **Next Generation Sequencing:** Library preparation was performed using the NEBNext DNA Ultra II library preparation kit for Illumina and performing 12 cycles PCR amplification with unique dual indexed primers. Libraries were pooled at equimolar ratios, and subject to 150 base paired end sequencing on an Illumina NovaSeq 6000 at Azenta Life Sciences. **Mass spectrometry:** Protein was on-bed digested with trypsin, without reduction or alkylation. Peptides were desalted by C18 ZipTip before drying and resuspending in aqueous 0.1% TFA for LC-MS. Peptides were acquired over a 1 h acquisition with elution from a 50 cm EN PepMap C18 column driven by a Waters mClass UPLC onto an Orbitrap Fusion Tribrid mass spectrometer operated in DDA mode. MS1 spectra were acquired in the Orbitrap mass analyser at 120K resolutions and a cycle time of 1 s, with MS2 spectra acquired in parallel in the linear ion trap using TopSpeed selection.

**Next Generation Sequencing Analysis:** Read quality was assessed with FastQC v0.11.7 [77], removing any retained adapters and poor quality reads using a sliding window approach with trimmomatic v0.36 [78]. Reads were discarded when trimmed to less than 50 base pairs and only intact read pairs were retained. Reads were aligned to the GRCm39 primary mouse assembly with STAR v2.7.10a [79] and annotated with SAMtools v1.15 [80] fixmate and markdup. ENCODE mm10 'blacklist' regions [81]) were removed after GRCm39 UCSC liftOver using BEDTools v2.30.0 [82]. Peaks were called with MACS v2.2.7.1 [83] and pairwise differential binding assessed using DiffBind v3.14.0 [84] in R v4.3.3 using all replicates and input controls. Absolute $log_2$ fold changes greater than 1 and FDR-corrected p values less than 0.05 were considered in the first instance, with FDR-based ranking also used to assess condition-specific HIF-1α binding importance. Relevant peak locations were plotted using karyoploteR v1.3.0 [85].

Peak locations were annotated using ENCODE and GENCODE GRCm39 annotations and HREs detected using Jellyfish 2.3.0 [86].

**Mass spectrometry Analysis:** LC-MS chromatograms in. raw format were imported into Progenesis QI for peak picking and chromatographic alignment, before exporting a concatenated tandem mass list in .mgf format for Mascot database searching. Searching was performed against the mouse subset of the SwissProt database appended with common proteomic contaminants. Peptide spectral matches were filtered to 1% FDR using the percolator algorithm and as assessed empirically against a reverse database search. Accepted peptide identifications were imported into Progenesis QI, associated with the chromatographic data and mapped between runs. Data were further filtered to require a minimum of two unique peptides per protein quantification. Relative protein quantification was inferred from peptide precursor ion areas using unique, non-conflicting peptides only and after normalisation to total peptide intensity among runs. Statistical testing between groups was done using anova with the null hypothesis being that proteins are of equal abundance between all samples. The Hochberg and Benjamini FDR methodology was used for multiple test correction of p-values to q-values.

**Patient data**

We obtained bulk RNA-sequencing data and clinical annotations from 230 patients with a positive JAK2V617F mutation from the Princess Margaret Cancer Centre MPN cohort [48]. Individual gene expression counts were subjected to VST normalisation using the 'DESeq2' R package whereas gene sets were scored using gene set variation analysis (GSVA) using the 'GSVA' R Package [50,74]. Survival analysis was performed using the 'survival' and 'survminer' packages in R and visualised with Kaplan Meier plots to depict differences in survival outcomes between patients with high or low gene expression or gene set enrichment. MIPSS70 analysis was performed within 172 patients with available genomic annotations [51].

**Gene signature identification**

We first identified the disparate genes between the VF_Hx/VF_Nx signatures (n=158), and the VF_Hx/WT_Hx signatures (n=144), representing those genes gained and lost, respectively, HIF-1 target genes in the VF_Hx context (Fig.7A). To delineate genes that are acquired as HIF-1 targets as a result of chromatin alterations and not aberrant HIF-1 function, we next screened out those with altered accessibility [52]. These genes were then analysed for differential expression in bulk and scRNAseq JVF vs WT [53,54], and finally differentially expressed genes were analysed for correlation with progression/survival and AML transformation in the JVF MPN patient cohort.

**Animal models**

Animals used in this study were C57BL/6 mice (WT) and Jak/2E/B6 Stella Cre heterozygous and homozygous. All mice were bred and housed under specific pathogen-free conditions at the University of York. All animal procedures were approved by the University of York Animal Welfare and Ethical Review Board and carried out under United Kingdom Home Office license (PEAD116C1). Animals were euthanized by $CO_2$ asphyxia and cervical dislocation. All the mice were 8 to 12 weeks old.

**Lineage negative HSPC isolation and expansion**

Lineage negative cells were isolated from the pelvis, femurs and tibias of C57BL/6 (WT) and Jak/2E/B6 Stella Cre mice using EasySep™ Mouse Hematopoietic Progenitor Cell Isolation Kit (STEMCELL Technologies 19856) according to the manufacturer's protocol, and were either used directly in downstream experiments or FACS sorted into Fibronectin-coated plates (Corning) containing PVA mouse expansion media [75]: Ham's F12 media (Gibco) supplemented with 1X penicillin-streptomycin-glutamine (Gibco), 1X HEPES (Gibco), 1X insulin-transferrin-selenium-ethanolamine (Gibco), 1 mg/ml polyvinyl alcohol (Sigma), 100 ng/ml recombinant mouse thrombopoietin (TPO; Peprotech), and 10 ng/ml recombinant mouse stem cell factor (SCF; Peprotech)1,2. 20 nM, IACS-010759 (Selleck). Gating strategy for FACS sorting of ESLAMS ($CD48^-$, $CD45^+$, $Sca1^+$, $CD150^+$, $EPCR^+$) HSCs as follows; FSC-A vs SSC-A plotted to gate cells and exclude debris, FSC-H vs FSC-A plotted to gate 'singlets' and exclude 'doublet' events, SSC-H vs SSC-A plotted to gate 'singlets' and exclude 'doublet' events, Live/Dead Violet (LIVE/DEAD™ Fixable Violet Dead Cell Stain (Invitrogen L34955)) vs FSC-A plotted to gate for live cells and exclude dead cells, CD150 (PE/Cyanine7 anti-mouse CD150 (SLAM) Antibody (Biolegend)) vs CD48 (APC anti-mouse CD48 Antibody (Biolegend)) plotted to gate $CD150^+$ $CD48^-$ (SLAM) cells, EPCR (CD201 (EPCR) Monoclonal Antibody (eBio1560 (1560)), PE, eBioscience™ (Invitrogen)) vs CD45 (Alexa Fluor® 488 anti-mouse CD45 Antibody (Biolegend)) plotted to gate for $EPCR^+$ $CD45^+$ (ESLAM) cells, EPCR vs Sca-1 (Brilliant Violet 605™ anti-mouse Ly-6A/E (Sca-1) Antibody (Biolegend)) plotted to gate for $Sca\text{-}1^+$ cells (ESLAMS). Endpoint immunophenotyping of expanded cultures was performed after 14 days by flow cytometry with panel of antibodies as previously described (Fig.M1) [76] including ViaKrome 808 Fixable Viability Dye (Beckman Coulter), CD34 - CD34 Monoclonal Antibody (RAM34) Alexa Fluor™ 488, eBioscience™ (Invitrogen), CD16/32 - PE/Cyanine7 anti-mouse CD16/32 Antibody (Biolegend), CD41 - APC anti-mouse CD41 Antibody (Biolegend), Lineage - PE anti-mouse Lineage Cocktail (Biolegend) (Fig.M1-M2 for gating strategy and exemplary flow plots).

For drug treatments of expansion cultures, 100 nM (or dose as on figure μM) GN44028 (MedChemExpress or Tocris-Biotechne), SMI-4a (Cayman Chemical) (dose as shown on figure μM), TCS7009 (Tocris-Biotechne) (dose as shown on figure μM) or DMSO (Sigma) were added to the HSC media where indicated. Cells were incubated at 5% $CO_2$ and either 1% $O_2$, 5% $O_2$ or 20% $O_2$ (Invivo 500 hypoxic workstation or Binder CO2 incubator) for 6 hours as previously described [7]. Re-plating assays were performed by FACS sorting cultured live (PI-) cells into fresh wells of complete media and then assessing the derived cell cultures after 14 days. We used >4000 live cells/well with >10% c-Kit+Sca1+Lineage- cells as the cut-off for replating activity. Extreme limiting dilution analysis was used to estimate culture replating cell frequencies.

**Flow cytometry.**

Following BM isolation and lineage depletion, cells were resuspended in HSC culture media (see above) and cultured at 5% $CO_2$ and either 1% $O_2$ or 20% $O_2$ (Invivo 500 hypoxic workstation or Binder CO2 incubator) for 4 hours as previously described [7]. Cells were then fixed (16% PFA (2% PFA final concentration)), permeabilised in methanol and stained with LIVE/DEAD™ Fixable Violet Dead Cell Stain (Invitrogen L34955) and either HIF-1α (D1S7W) XP® rabbit mAb (Alexa Fluor® 488 conjugate) (Cell Signalling Technologies) or HIF-1 alpha Monoclonal Antibody (Mgc3), PE, eBioscience™ (Invitrogen) with appropriate isotype control (Rabbit (DA1E) mAb IgG XP® Isotype Control (Alexa Fluor® 488 conjugate) (Cell Signalling Technologies) or Mouse IgG1 kappa Isotype Control (P3.6.2.8.1), PE, eBioscience™ (Invitrogen)). Samples were washed with FACS buffer and run on CytoFlex LX flow cytometer (Beckman Coulter) and data analysed using FlowJo software (BD).

**Multiplex fluorescent barcoding and phospho-flow cytometry.**

UT7/TPO cells were cultured as indicated and then starved overnight in 2% FBS RPMI (Roswell Park Memorial Institute (RPMI) 1640 Medium (Gibco)) media with no TPO prior to treatment. Cells were treated as indicated with 2.5μm SMI-4a (Cayman Chemical) for 6hrs, 2μm Ruxolitinib (Rux) (Cell Guidance Systems) for 6hrs, DMSO (Sigma) vehicle control for 6hrs and/or 50ng/ml TPO (Peprotech) 30 minutes in either 6hrs of hypoxic (1% $O_2$) conditions or in normoxia (20% $O_2$). Cells were then fixed in 16% PFA (16% Formaldehyde, Methanol-Free #12606 Cell Signalling Technology) (2% PFA final concentration), resuspended in methanol for permeabilization and stored at -80℃ prior to barcoding. Compensation controls prepared prior to barcoding using cells for dye single stains and VersaComp beads (Beckman Coulter B22804) for signalling antibodies' single stains. Barcoding and flow cytometry was performed as previously described (including antibodies specified), [32]. Fig.S6 for gating strategy.

**Cell culture**

Murine BaF3 cells stably expressing the human MPL and human JAK2 (BaF3 hMPL hJAK2 WT) or the JAK2V617F mutant (BaF3 hMPL hJAK2 VF) were grown in RPMI media (Gibco) (10% FBS (Gibco), 1% penicillin (Gibco)/streptomycin/L-glutamine (Gibco)) (plus murine IL-3 (Peprotech) for hJAK2 WT cells). Human UT7/TPO cells expressing JAK2 (UT7/TPO JAK2 WT) or the JAK2V617F mutant (UT7/TPO JAK2 VF) were grown in RPMI media (Gibco) (10% FBS (Gibco), 1% penicillin (Gibco)/streptomycin/L-glutamine

(Gibco)) (plus human TPO (Peprotech) for JAK2 WT cells). Human SET2 cells heterozygous for the JAK2V617F mutation were grown in RPMI media (Gibco) (20% FBS (Gibco), 1% penicillin/streptomycin (Gibco)/L-glutamine (Gibco)) and human HEL cells homozygous for the JAK2V617F mutation were grown in RPMI media (Gibco) (10% FBS (Gibco), 1% penicillin/streptomycin (Gibco)/L-glutamine (Gibco)). Cells were incubated at 5% $CO_2$ and the indicated $O_2$ concentrations.

**Western blot**

Cells were cultured as above and treated with either 4mM NAC (Biotechne), 75μM DMF (Stratech Scientific), 10μM SMI-4a (Cayman Chemical), (dose as on figure μM) TCS PIM1.1 (Santa Cruz Biotechnology), (dose as on figure μM) PIM447 (Sellachem), 2μM Ruxolitinib (Rux) (Cell Guidance Systems), 10μM MG132 (MedChemExpress), 50ng/ml human TPO (Peprotech) or (dose as on figure nM) SCH874 (Sellachem) as indicated. Cells were lysed in cell lysis buffer (20 mM Tris-HCl (pH 7.5), 150 mM NaCl, 1 mM $Na_2EDTA$, 1 mM EGTA, 1% Triton) plus Halt™ Protease and Phosphatase Inhibitor Cocktail, EDTA-free (ThermoFisher Scientific). Lysate (20 μg) was separated on 4-20% Mini-PROTEAN® TGX Stain-Free™ protein gels (Biorad) and transferred onto Immun-Blot® Low Fluorescence PVDF membrane (Biorad). Membranes were probed for the indicated antibodies (list below).

Isotype control - Rabbit (DA1E) mAb IgG XP® isotype control (Cell Signalling Technology)

HIF-1α - HIF-1α (D1S7W) XP® Rabbit mAb (Cell Signalling Technology)

pHIF-1α (Thr455) - Rabbit mAb (Generously donated by Prof. Noel Warfel)

β-Actin - Anti-actin hFAB Rhodamine (Biorad)

TDP-43 - TARDBP Antibody (H-8): sc-376532 mouse mAb (Santa Cruz)

ERK1/2 - p44/42 MAPK (Erk1/2) (L34F12) mouse mAb #4696 (Cell Signalling Technologies)

pERK1/2 - Phospho-p44/42 MAPK (Erk1) (Tyr204)/(Erk2) (Tyr187) (D1H6G) Mouse mAb #5726 (Cell Signalling Technologies)

Golga3 - Golgin 160 Antibody (C-8): sc-374596 mouse mAb (Santa Cruz)

SIK2 - SIK2 Antibody (B-12): sc-393139 mouse mAb (Santa Cruz)

IMPDH -IMPDH Antibody (F-6): sc-166551 mouse mAb (Santa Cruz)

p70 S6K - p70 S6 kinase α Antibody (H-9): sc-8418 mouse mAb (Santa Cruz)

pp70 S6K (T389) - Phospho-p70 S6 Kinase (Thr389) (108D2) Rabbit mAb #9234 (Cell Signalling Technologies)

PIM1 - PIM1 Recombinant Polyclonal Antibody (19HCLC) (Invitrogen)

NRF2 - Nrf2 Antibody (A-10): sc-365949 mouse mAb (Santa Cruz)

STAT3 - STAT3 (124H6) mouse mAb (91395) (Cell Signalling Technologies)

pSTAT3 (Y705) - Phospho-STAT3 (Tyr705) (D3A7) XP® rabbit mAb (91455) (Cell Signalling Technologies)

JAK2 - JAK2 Monoclonal Antibody (691R5) mouse mAb (Invitrogen)

pJAK2 (Y1007/8) - Phospho-Jak2 (Tyr1007/1008) (C80C3) Rabbit mAb #3776 (Cell Signalling Technologies)

**Immunoprecipitation**

Cells were cultured as above with the indicated conditions. Cells were lysed in cell lysis buffer (20 mM Tris-HCl (pH 7.5), 150 mM NaCl, 1 mM $Na_2EDTA$, 1 mM EGTA, 1% Triton) plus Halt™ Protease and Phosphatase Inhibitor Cocktail, EDTA-free (ThermoFisher Scientific). 200 µg of cell lysate used as input for immunoprecipitation. Cell lysates were precleared by addition of Protein A magnetic beads (Cell Signaling Technologies) for 20 minutes at RT with rotation followed by magnetic separation. Proteins were captured by addition of HIF-1α (D1S7W) XP® Rabbit mAb (Cell Signaling Technology) or TDP-43 sc-376532 mouse mAb (Santa Cruz) and incubated overnight at 4°C with rotation followed by immunoprecipitation by addition of Protein A magnetic beads and incubation for 20 minutes at RT, followed by magnetic separation and washing. Immunoprecipitated protein as then analysed by western blotting.

**Phosphoproteomics**

Samples immunoprecipitated for HIF-1α (see above) were digested on-bead using the RIME protocol and trypsin as the protease, before $TiO_2$ (MagReSyn) phosphopeptide enrichment. Resulting peptides were loaded onto an EvoTip Pure for introduction onto an 8 cm performance column. Elution was carried out using a 60 SPD pre-set gradient on an EvoSep One UPLC. DDA-PASEF data were acquired using a Bruker timsToF HT. Data were searched through FragPipe for phosphopeptide identification and relative quantification at 1% FDR. Statistical testing was performed for all pairwise comparisons using limma via FragPipe-Analyst. Zero value imputation was applied and the Hochberg and Benjamini approach was used for multiple test correction. Significance threshold was set at $q < 0.05$.

**Bulk RNAseq and analysis**

Bulk RNAseq data of JAK2V617F knock-in mice re-analysed from publicly available dataset (GEO: GSE180853). RNA extracted, prepared and sequenced as per method specified in paper [53]. Briefly, RNA extracted by TRIzol extraction from sorted-cell populations as indicated. RNA-seq libraries were generated by 3' sequencing and SMART-Seq2 amplification and sequenced on an Illumina HiSeq 4000. SRA short read

RNA-seq analysis performed as follows; raw, paired-end Illumina reads downloaded from NCBI SRA were first trimmed with Cutadapt (version 2.10) [87] to remove the Illumina universal adapters and any low-quality sequence, with a minimal length threshold of 20bp. Trimmed reads were mapped and quantified using the Salmon (version 1.10) [88] pseudo-aligner, using the GENCODE M32 (GRCm39) [89] Mus musculus transcriptome, specifying the ISR library type, 100 bootstraps, and the "validateMappings" option. Differential expression analysis was run using the Sleuth (version 0.30.1) [90] and Wasabi (version 1.0.1) [91] packages in R, on a gene-wise basis, and using the Wald test to find statistically significant, differentially expressed genes. Genes with a log2 fold change of greater than +/-1 and with a q value of less than 0.1 were taken to be significantly differentially expressed.

**scRNAseq and analysis**

scRNAseq of MPN patients and healthy controls taken from publicly available dataset (GEO: GSE144568). RNA extracted, prepared and sequenced as per method specified in paper [54]. SRA single cell RNA-seq analysis performed as follows; CellRanger files downloaded from NCBI SRA were loaded into the Seurat package (version 5.1.0) [92] and the Read10X function. Samples read in separately and were filtered to exclude genes expressed in fewer than 3 cells, and to exclude cells with fewer than 500 genes expressed. Cells with greater than 10% mitochondrial gene expression were excluded. Filtered samples were merged and then normalised, scaled, and variable features identified. Layers in the merged object were integrated with the "RPCAIntegration" method. The "FindNeighbors" function was run using the first 30 principal components, followed by FindClusters function with a resolution of 0.25. The RunUMAP function was used to calculate the UMAP dimensional reduction, again with the first 30 principal components. Marker genes for the UMAP clusters were found by first joining the layers with the JoinLayers function and then using the FindAllMarkers function. The UMAP clusters were renamed according to their cell types, by manually assessing the marker genes for each cluster. HIF gene module scores were calculated by providing a list of HIF-associated genes to the R UCell package [93] and the "AddModuleScore_UCell" function, and added to the Seurat object to visualise the HIF gene module scores for the cells in their respective UMAP clusters.

**RT-qPCR**

RNA was extracted from samples using Monarch® Total RNA Miniprep Kit #T2010S and RT performed using LunaScript RT Master Mix Kit (Primer-free) #E3025S (New England Biolabs) as per manufacturer protocols with SimpliAmp™ Thermal Cycler (Applied Biosystems™ #A24811). qPCR was performed with SYBR™ Green PCR Master Mix (Applied Biosystems™ #4309155) and primers: Ms GAPDH (F): CATCACTGCCACCCAGAAGACTG, Ms GAPDH (R): ATGCCAGTGAGCTTCCCGTTCAG, Ms PIM-1 (F): CGCGACATCAAGGACGAGAACA, Ms PIM-1 (R):

CGAATCCACTCTGGAGGACTGT on StepOnePlus™ Real-Time PCR System (Applied Biosystems™ #4376600). Analysis performed on StepOne Software.

Fig. M1

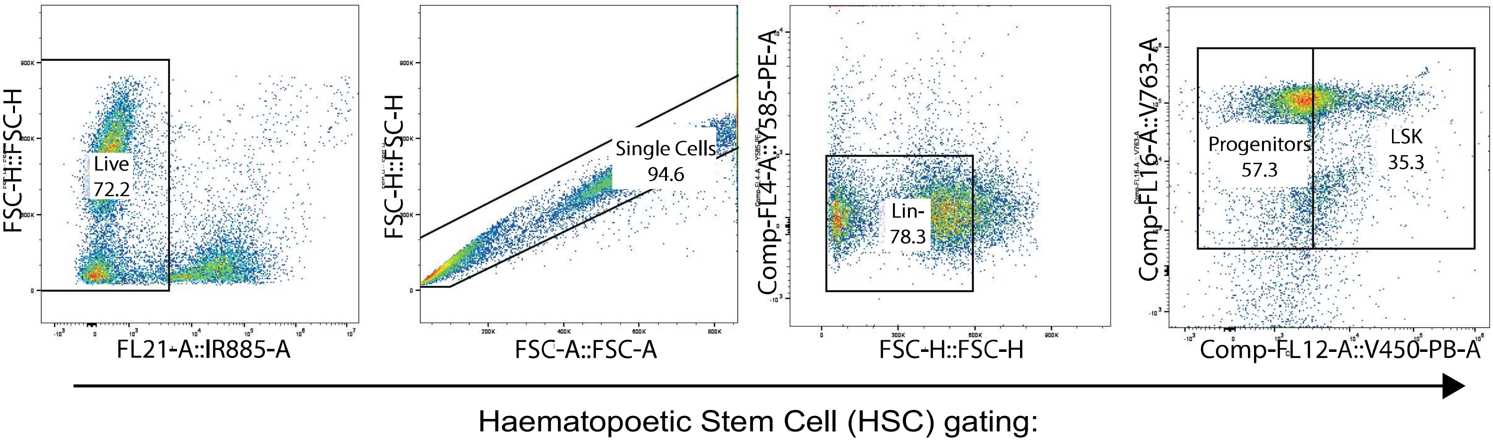
FSC-H::FSC-H
Live
72.2
FL21-A::IR885-A
FSC-H::FSC-H
Single Cells
94.6
FSC-A::FSC-A
Comp-FL4-A::Y585-PE-A
Lin-
78.3
FSC-H::FSC-H
Comp-FL16-A::V763-A
Progenitors
57.3
LSK
35.3
Comp-FL12-A::V450-PB-A
Haematopoetic Stem Cell (HSC) gating:

Fig. M2

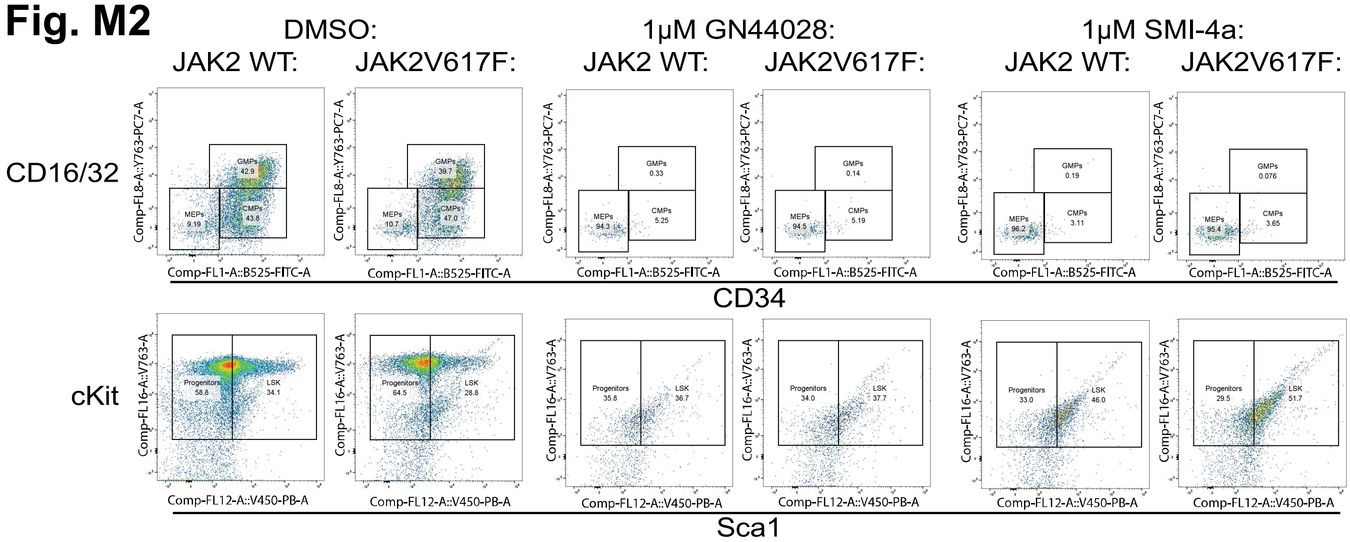
DMSO:
1μM GN44028:
1μM SMI-4a:
JAK2 WT:
JAK2V617F:
JAK2 WT:
JAK2V617F:
JAK2 WT:
JAK2V617F:
CD16/32
Comp-FL8-A::Y763-PC7-A
GMPs 42.9
MEPs 9.19
CMPs 43.8
GMPs 39.7
MEPs 10.7
CMPs 47.0
GMPs 0.33
MEPs 94.3
CMPs 5.25
GMPs 0.14
MEPs 94.5
CMPs 5.19
GMPs 0.19
MEPs 96.2
CMPs 3.11
GMPs 0.076
MEPs 95.4
CMPs 3.65
Comp-FL1-A::B525-FITC-A
CD34
cKit
Comp-FL16-A::V763-A
Progenitors 58.8
LSK 34.1
Progenitors 64.5
LSK 28.8
Progenitors 35.8
LSK 36.7
Progenitors 34.0
LSK 37.7
Progenitors 33.0
LSK 46.0
Progenitors 29.5
LSK 51.7
Comp-FL12-A::V450-PB-A
Sca1

**Supplementary Figure Legends:**

S1(A) Cell count data for Fig.1D

(B) Violin plot of flow cytometry data from Fig.1D normalised to isotype.

(C) Immunoblot for HIF-1α in primary mouse lineage depleted HSCs from wildtype, heterozygous and homozygous JVF mice cultured in 1% O2 hypoxic conditions for the indicated length of time. HIF-1α antibody used with β-actin as loading control (n=2). Quantification by densitometry (below).

S2 Replating efficiency from cultures described in Fig.1(C).

S3(A) HIF-1 inhibition partially rescues HSC expansion culture phenotype in JVF cells as determined by proportion of LSK cells in culture, returning the LSK cell percentage closer to those observed in vehicle treated wildtype expansion cultures. In wildtype cultures however, HIF-1 inhibition decreased LSK percentage. Ex-vivo expansion of long-term HSCs from wildtype (purple) and JVF heterozygous (blue) mice in the PVA culture system treated with the indicated dose (μM) of GN44028 or vehicle control and immunophenotyped (by gating strategy following previously described HSC populations [76]) after 14 days of expansion and expressed as percentage of lineage negative cell population (n=3).

(B) HIF-1 inhibition partially rescues HSC expansion culture phenotype in JVF cells as determined by proportion of progenitor cells in culture, returning the progenitor cell percentage to those observed in vehicle treated wildtype expansion cultures. In wildtype cultures, HIF-1 inhibition leaves progenitor cell percentage unaffected.

S4(A) Confirmatory representative western blots of Fig.1C NAC ROS inhibition assay in SET2 and HEL cell lines.

(B) Treatment with ROS scavenger NAC (N-acetyl-L-cysteine) fails to inhibit JVF-induced HIF-1α stabilisation, whereas treatment with the ROS inhibitor DMF (dimethyl fumarate) does inhibit JVF-induced HIF-1α stabilisation. Representative immunoblot of HIF-1α, PIM1 and DMF-target NRF2 levels in wildtype and JVF UT7/TPO cells, SET2 cells and HEL cells treated as indicated with either NAC, DMF or vehicle control. Immunoblotting carried out using HIF-1α, NRF2 and PIM1 antibodies with β-actin as a loading control (n=3).

S5 Treatment with ERK inhibitor SCH874 fails to destabilise JVF-induced HIF-1α stabilisation. Immunoblot of HIF-1α, pJAK2, ERK1/2 and pERK1/2 in JVF UT7/TPO cells, treated as indicated with either SCH874, Rux or vehicle control. Immunoblotting carried out using HIF-1α, pJAK2 Y1007/Y1008, pERK1/2 Y204/Y187 and ERK1/2 antibodies with β-actin as a loading control (n=2).

S6 Gating strategy for multiplex fluorescent barcoding and phospho-flow cytometry.

S7 PIM1 transcript expression is higher in JVF mouse LSKs compared to wildtype. Bulk RNA-seq analysis of mouse LSKs from wildtype (n=3) and JVF heterozygous mice (n=3).

S8 Confirmatory representative western blot of Fig.2A(I) PIM1 inhibition assay in wildtype UT7/TPO cells (n=3).

S9 Confirmatory representative western blot of Fig.2A(I) PIM1 inhibition assay in wildtype and JVF hMPL BaF3 cells (n=3).

S10 Treatment with SMI-4a effectively leaves JAK2 signalling cascades untouched, whilst treatment with Rux ablates JAK2 downstream signalling. Barcoded phospho-flow cytometry of kinase cascade constituent proteins downstream of JAK2 in JVF and wildtype UT7/TPO cells, cultured in normoxic (20% O2) or hypoxic (1% O2) conditions, starved overnight or treated with TPO and treated with SMI-4a, Rux or vehicle control as indicated. Heatmap of mean channel fluorescence intensity (MFI), data normalised based on fold change to mean value of wildtype starved, no TPO, normoxic condition (±SEM) (n=3).

S11 HIF-2 inhibition has no significant effect on expansion culture phenotype as determined by proportion of LSK/progenitor cells in culture in either wildtype or JVF cells. Ex vivo expansion of long-term HSCs from wildtype (purple) and JVF heterozygous (blue) mice in the PVA culture system treated with the indicated dose (μM) of TCS7009 or vehicle control and immunophenotyped after 14 days of expansion and expressed as percentage of lineage negative cell population (n=3).

S12 Ser500 site is highly evolutionarily conserved across species whereas Thr498 is not. Sequence alignment of various species at Thr498/Ser500 site highlighted by blue dotted box as shown.

S13 Novel JVF-exclusive phosphorylation sites (Thr498 and Ser500) fall in HIF-1αs ODD (oxygen-dependent degradation) domain between canonical HIF-1α hydroxylation sites (P402 and P564). Ribbon diagram of HIF-1α, coloured by domain. Amino acid residues of interest marked in red.

S14 Levels of HIF-1α at 6hrs of hypoxic exposure are equal between wildtype and JVF hMPL BaF3 cells. Immunoblot of HIF-1α in wildtype and JVF hMPL BaF3 cells cultured in hypoxia for the indicated number of hours, blotted for HIF-1α with β-actin as loading control. Quantification by densitometry included.

S15(A) Full STRING functional cluster analysis of all cofactor proteins across all conditions identified by RIME-mass spec from Fig.4D.

(B) STRING functional cluster analysis of cofactors de-enriched (geomean relative to WT_Hx 0.01<0.005) for VF_Hx HIF-1α (left) and VF_Nx HIF-1α (right).

S16 Full panel of phosphorylated proteins identified by phospho mass spec as immunoprecipitated with HIF-1α in JVF and wildtype hypoxic UT7/TPO cells from Fig.3A(I) (n=3).

S17 Analysis of WT_Hx hMPL BaF3 cells by ChIP-seq shows canonical HIF signature gene loci binding. Volcano plot of loci binding intensity in WT_Hx cells relative to WT_Nx plotted by statistical significance (1/log10(BH-p)) with canonical HIF-1α target genes indicated by a cross and key signature genes and highly expressed non-signature genes labelled.

S18 Key HRE genes found to be differentially bound by HIF-1α in WT_Hx compared to WT_Nx include Vegfa and Slc2a3. Gene tracks of genomic loci where significant gain of HIF-1α binding as determined by ChIP-seq can be observed in WT_Hx (blue) compared to WT_Nx (purple).

S19 Weaker overall binding of HIF-1α to chromatin observed in normoxic and hypoxic JVF hMPL BaF3 cells compared to hypoxic-treated wildtype cells. Overlay of median genome-wide ChIP-seq binding signal at genomic regions centred at peaks of binding signals, in WT_Nx (purple), WT_Hx (blue), VF_Nx (yellow) and VF_Hx (green).

S20 JVF-stabilised HIF-1α binds to fewer genomic loci than wildtype hypoxia-stabilised HIF-1α, with no novel loci binding sites detected in JVF cells in either normoxia or hypoxia. Venn diagram of HIF-1α loci bound in the indicated conditions with number of genes unique to each condition labelled. Binding sites determined by ChIPseq in hMPL BaF3 cells (n=3).

S21 Across the mouse genome, HIF-1α binding appears lessened between chromosomes 1-4 and sites at chromosomes 8 and 14 appear prioritised in VF_Nx when compared to WT_Hx whilst the priority of HIF-1α binding appears enhanced in the region of chromosome 12 when compared to WT_Hx. Binding intensity of HIF-1α-bound genomic loci for VF_Nx (light blue) VF_Hx (dark blue) denoting bound more intensely by HIF-1α in JVF cells than WT_Hx, as determined by ChIP-seq, plotted by genomic location (chromosome).

S22 Cross analysis of mouse bulk RNA-seq data with JVF MPN patient scATAC-Seq data demonstrates no correlation between gene loci accessibility and gene transcription in JVF heterozygous mouse cells compared to wildtype. Plot of genome-wide gene accessibility (as derived from Izzo et al. Supplementary Figure 4 'Differential gene accessibility score in HSC and HSCMY clusters in untreated patients with myelofibrosis.') in primary JVF-mutation carrying patient cells plotted against gene expression (as determined by bulk RNA-seq of JVF heterozygous mice from publicly available dataset).

S23 A limited number of genes from canonical HIF-1 metagene signature (Lombardi et al.) identified in VF_Nx and VF_Hx gene signatures.

S24 Analysis of scATAC-seq data shows key VF_Hx 'signature' genes as more or less accessible in JVF-mutation carrying patient cells. Volcano plot of Log2FC (Fold Change) gene accessibility in JVF-carrying patients compared to healthy controls as determined by scATAC-seq, plotted against statistical significance (-Log10 FDR (False discovery rate)). Data derived from Izzo et al. Supplementary Figure 4 'Differential gene accessibility score in HSC and HSCMY clusters in untreated patients with myelofibrosis.'

S25 Log2 Fold Change expression of key genes with high expression in JVF cells, association with worse MPN patient disease severity and poorer survival outcomes from bulk RNA-seq analysis of JVF hypoxic cells compared to wildtype hypoxic mouse MEP and LSKs (n=3).

S26 BCL2L1 (BCL2XL) is significantly upregulated in transcript expression in MEPs/MkP and significantly associated with disease progression and survival. UMAP scRNAseq plot of control (top) and myelofibrosis (bottom) HSPCs with transcript of gene of interest (BCL2L1) highlighted in purple. The correlation of the individual genes of interest with MIPSS70 MPN patient classification and patient survival displayed below.

S27 TNKS, HECTD1, WIPF1, PPID are globally upregulated throughout HSPC clusters but were not significantly associated with disease progression and survival. UMAP scRNAseq plot of control (top) and myelofibrosis (bottom) HSPCs with transcript of genes of interest (TNKS, HECTD1, WIPF1, PPID) highlighted in purple. The correlation of the individual genes of interest with MIPSS70 MPN patient classification and patient survival displayed below.

S28 IER3 is significantly upregulated in more primitive progenitor populations and is inversely correlated with MPN patient disease severity and survival. UMAP scRNAseq plot of control (top) and myelofibrosis (bottom) HSPCs with transcript of genes of interest (IER3) highlighted in purple. The correlation of the individual genes of interest with MIPSS70 MPN patient classification and patient survival displayed below.

S29 No significant difference in expression of the VF_Hx signature genes as a whole can be observed in the 11 MPN patients from the JVF MPN cohort who spontaneously converted to AML (blue) compared to those that didn't (red).

S30 Graphical abstract/illustration of JVF-mediated HIF-1α stabilisation.

### Fig. S1

## A)

| Sample Name | Subset Name | Count | Mean : Comp-FL1-H | SD : Comp-FL1-H | Mode : Comp-FL1-H |
|---|---|---|---|---|---|
| 01-Hom 4h 1% O2-D6.fcs | Live | 121088 | 47505 | 22840 | 36286 |
| 01-Het 4h 1% O2-D4.fcs | Live | 42195 | 48894 | 20804 | 42805 |
| 01-WT 4h 1% O2-D2.fcs | Live | 32855 | 46162 | 15640 | 52771 |
| 01-Mixed Isotype-B4.fcs | Live | 45774 | 28721 | 10183 | 28476 |

| Sample Name | Subset Name | Count | Mean : Comp-FL1-H | SD : Comp-FL1-H | Mode : Comp-FL1-H |
|---|---|---|---|---|---|
| 01-WT 4h 1% O2-D2.fcs | Live | 32855 | 46162 | 15640 | 52771 |
| 01-WT 0h-B6.fcs | Live | 49560 | 36641 | 12102 | 36688 |
| 01-Mixed Isotype-B4.fcs | Live | 45774 | 28721 | 10183 | 28476 |

| Sample Name | Subset Name | Count | Mean : Comp-FL1-H | SD : Comp-FL1-H | Mode : Comp-FL1-H |
|---|---|---|---|---|---|
| 01-Het 4h 1% O2-D4.fcs | Live | 42195 | 48894 | 20804 | 42805 |
| 01-Het 0h-B8.fcs | Live | 54658 | 32025 | 11389 | 27249 |
| 01-Mixed Isotype-B4.fcs | Live | 45774 | 28721 | 10183 | 28476 |

| Sample Name | Subset Name | Count | Mean : Comp-FL1-H | SD : Comp-FL1-H | Mode : Comp-FL1-H |
|---|---|---|---|---|---|
| 01-Hom 4h 1% O2-D6.fcs | Live | 121088 | 47505 | 22840 | 36286 |
| 01-Hom 0h-B10.fcs | Live | 86879 | 26601 | 9811 | 25227 |
| 01-Mixed Isotype-B4.fcs | Live | 45774 | 28721 | 10183 | 28476 |

## B)

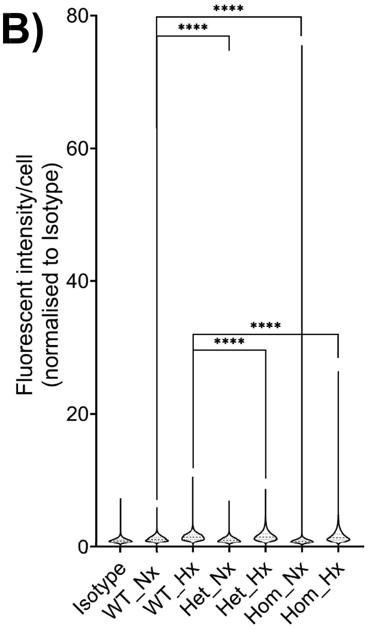


## C)

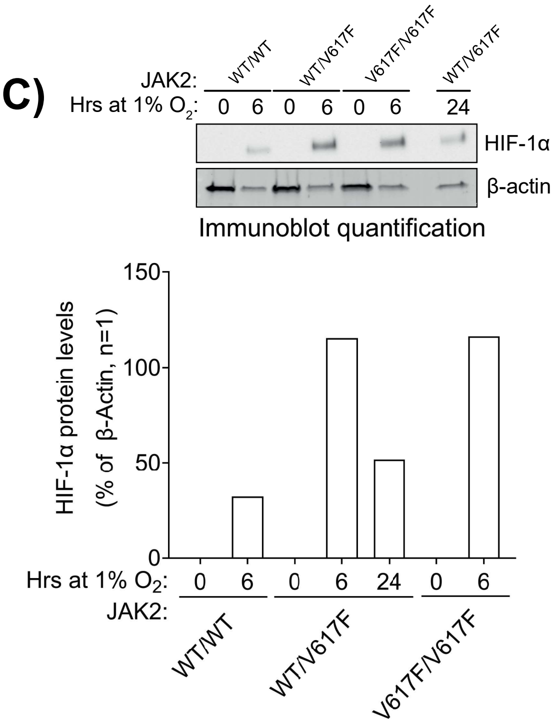


### Fig. S2

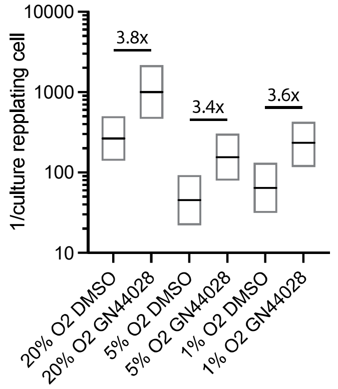


### Fig. S3

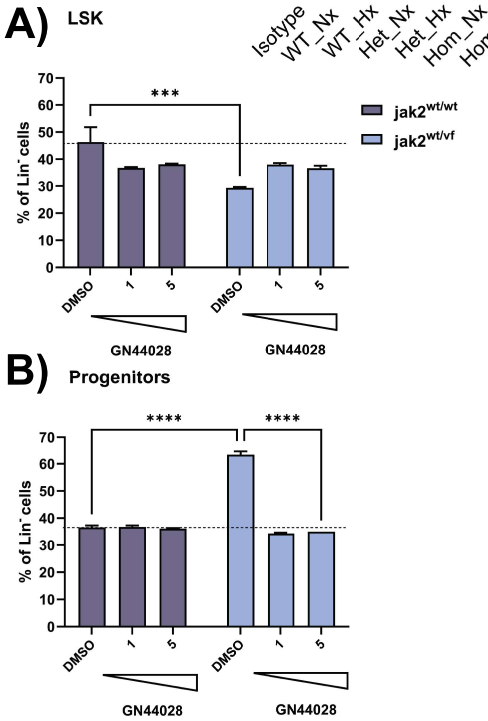


### Fig. S4

## A)

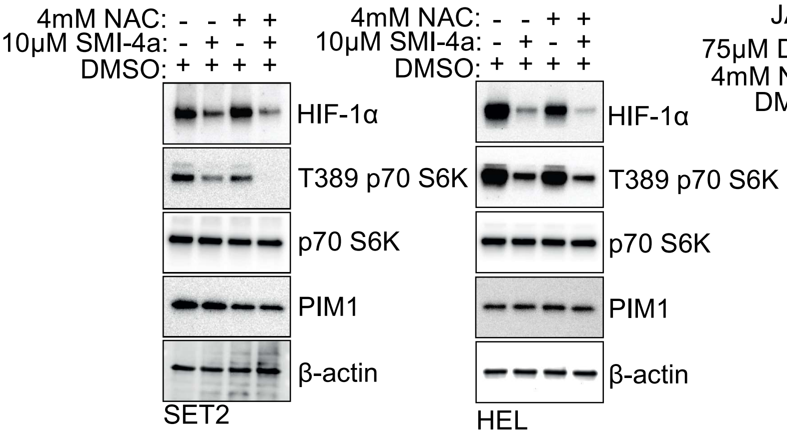


## B)

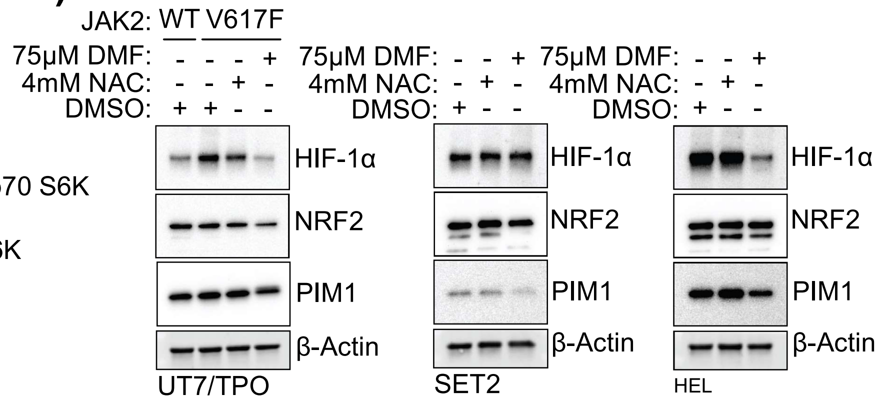

#### Fig. S5

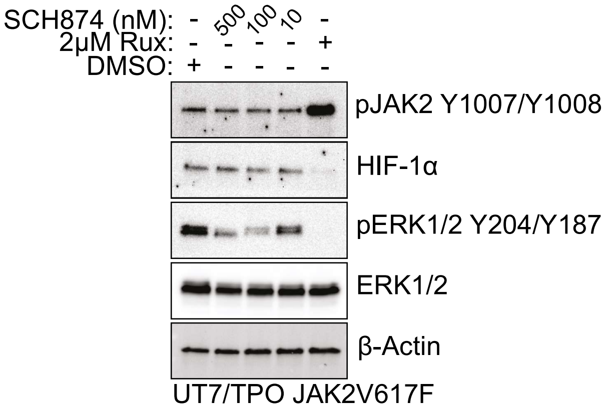


#### Fig. S6

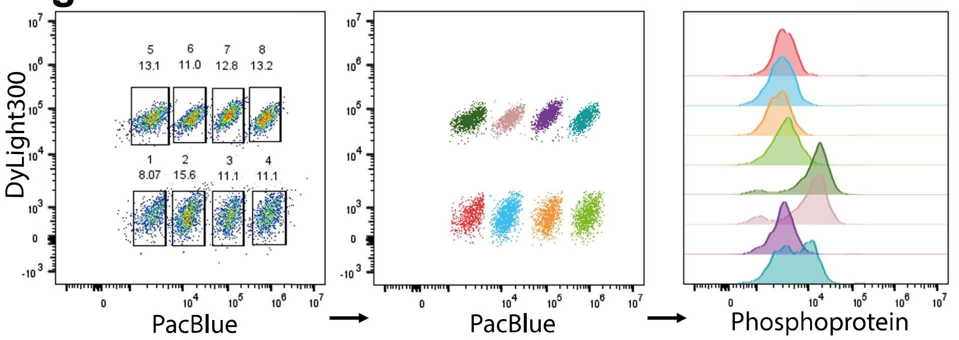


#### Fig. S7

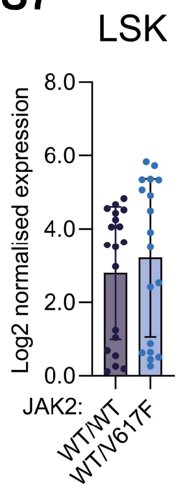


#### Fig. S8

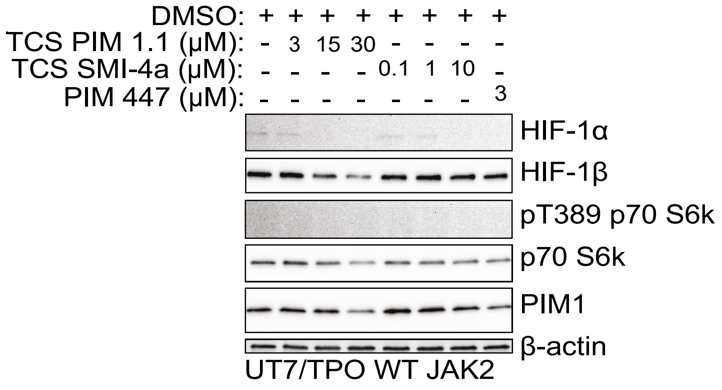


#### Fig. S9

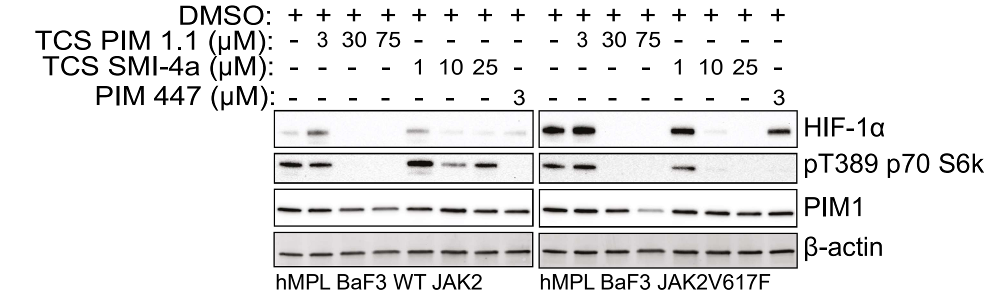


#### Fig. S10

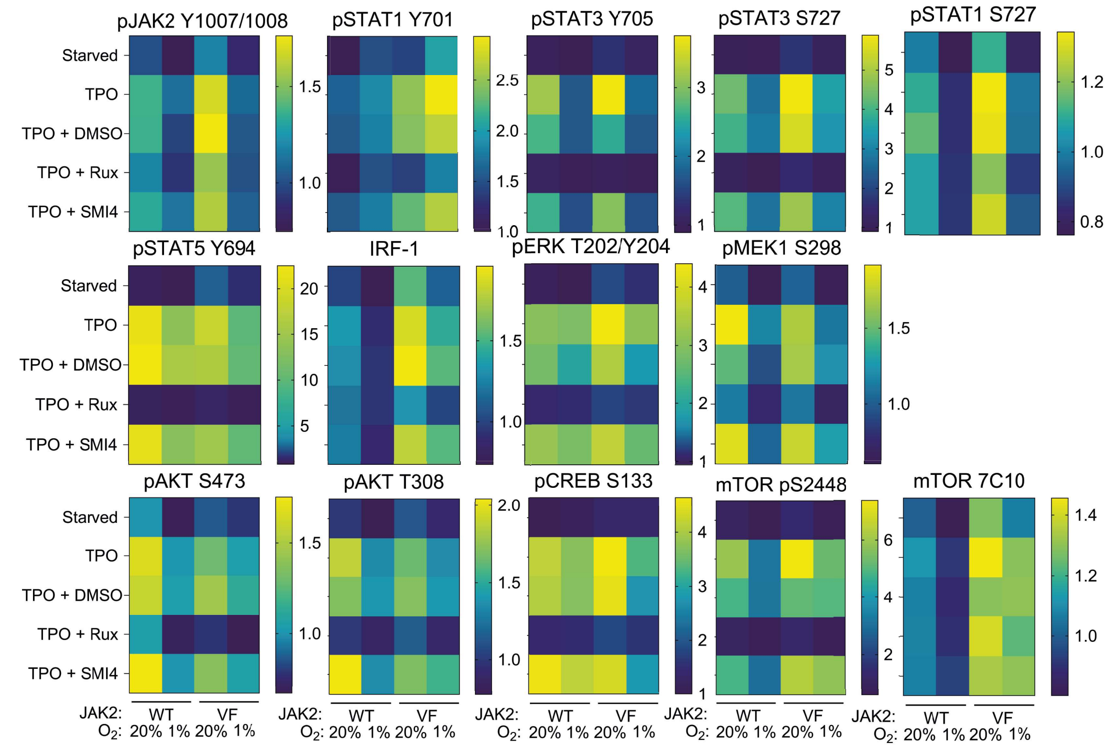

### Fig. S11

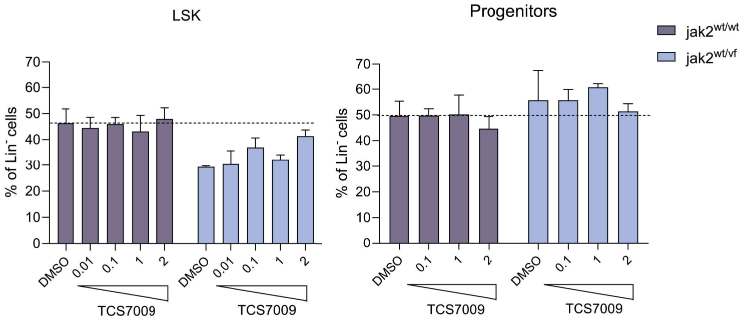


### Fig. S12

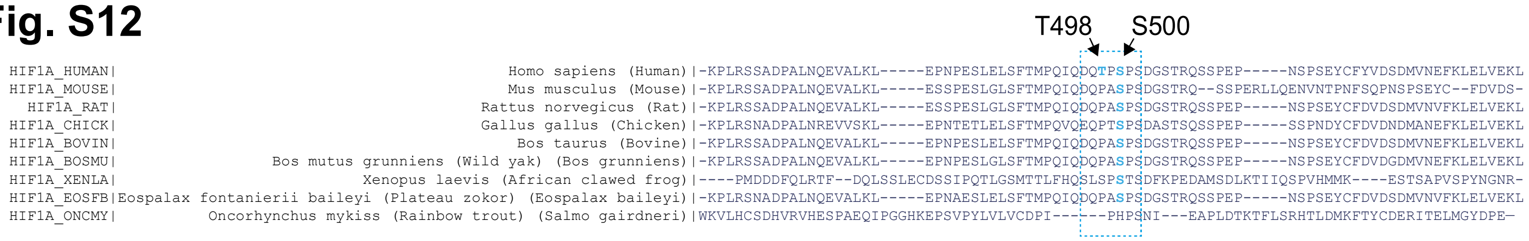


### Fig. S13

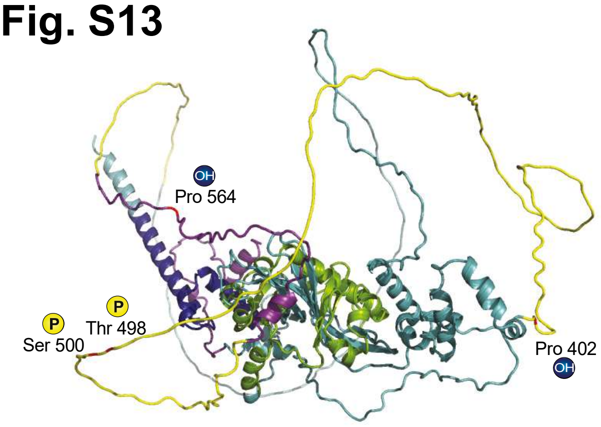


### Fig. S14

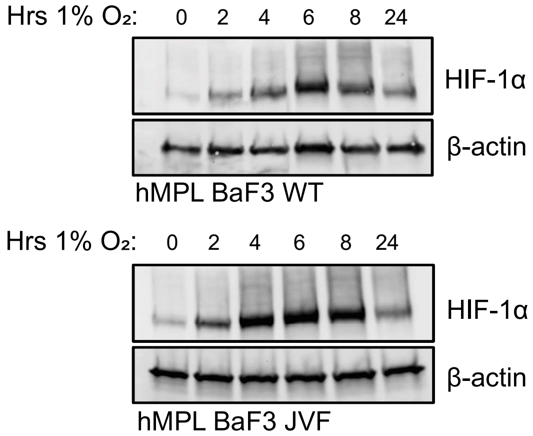


### Fig. S15

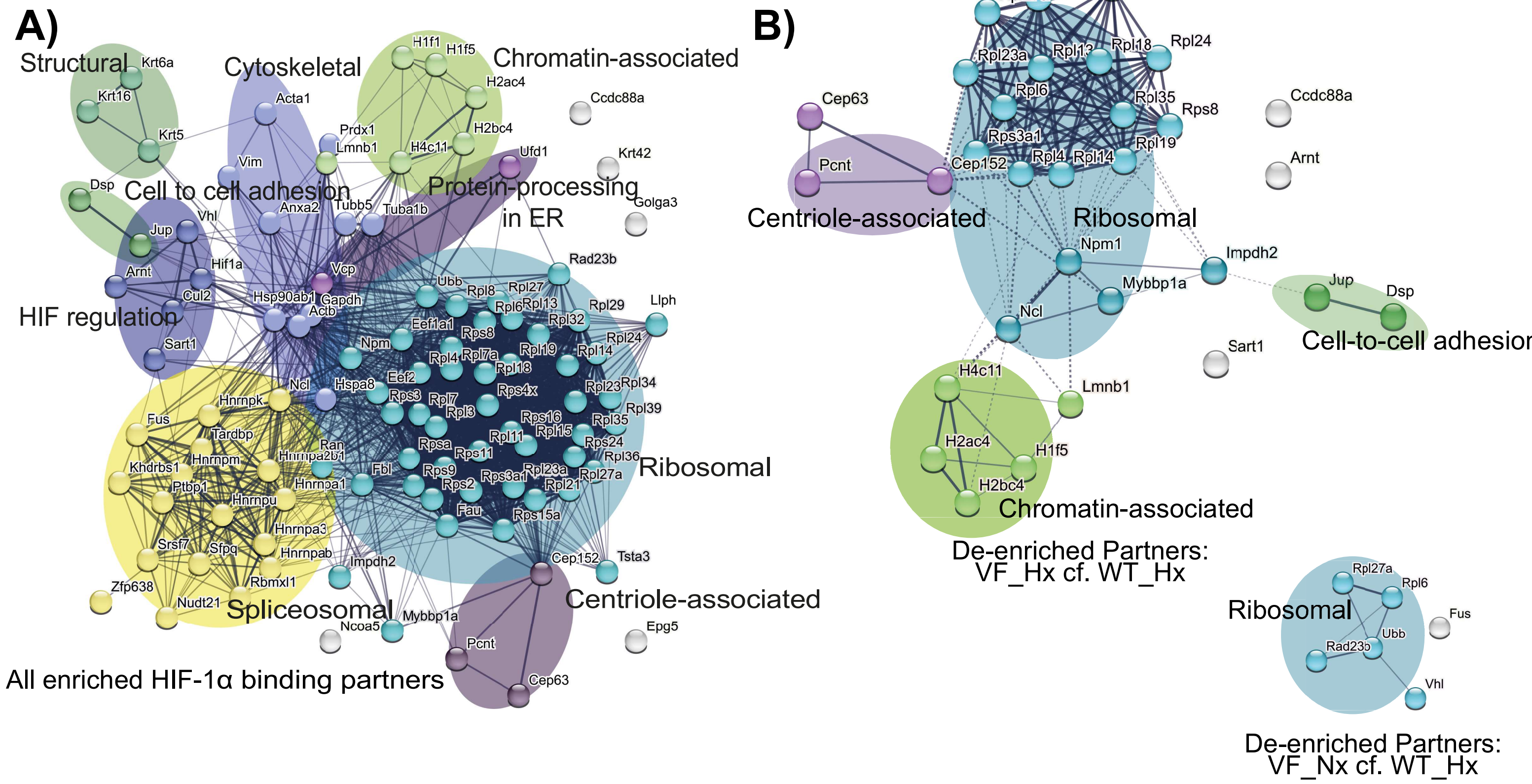

Fig. S16

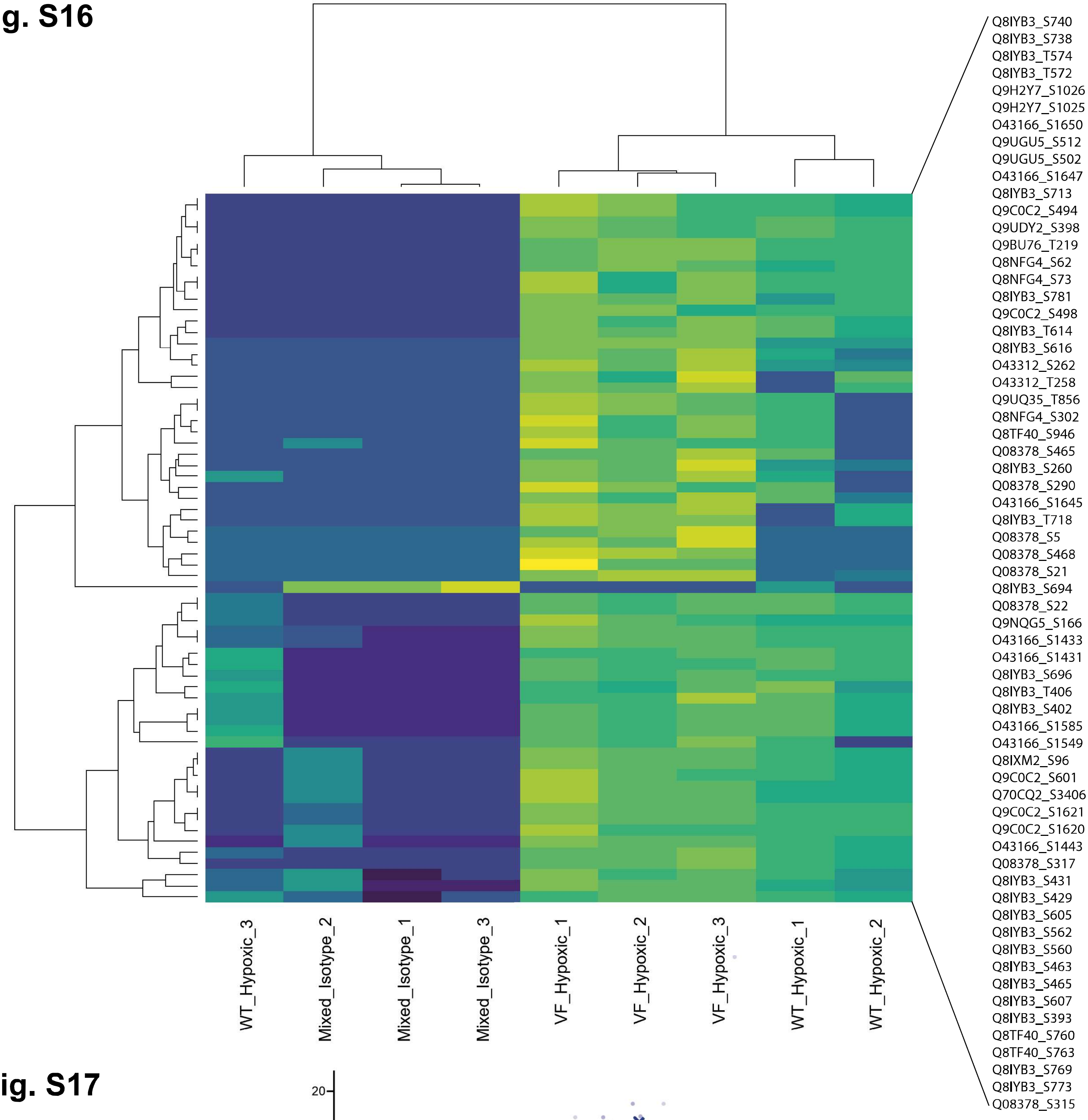


Fig. S17

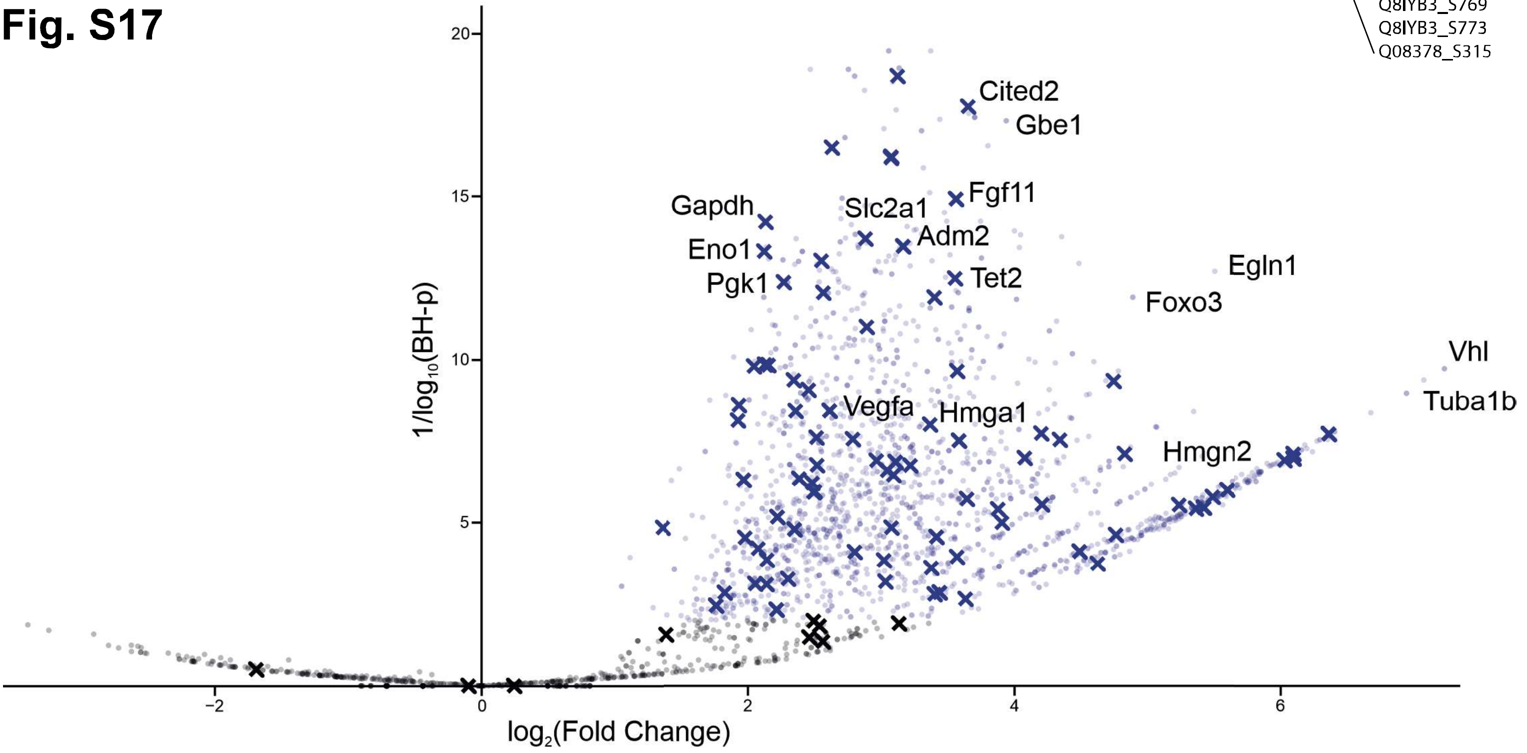

### Fig. S18

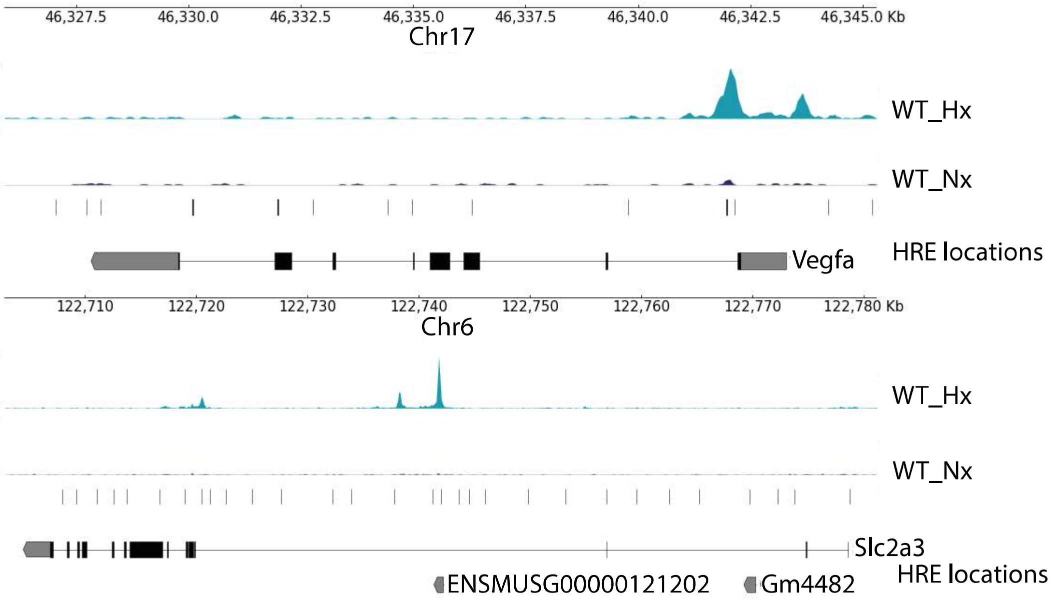


### Fig. S19

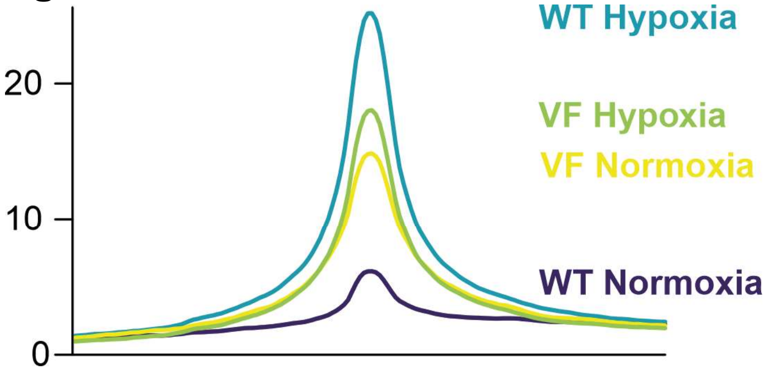


### Fig. S20

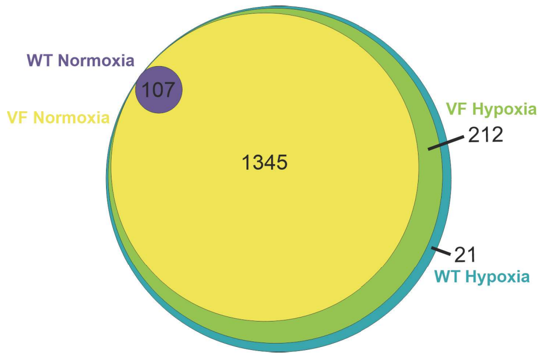


### Fig. S21

Binding intensity vs. WT_Hx

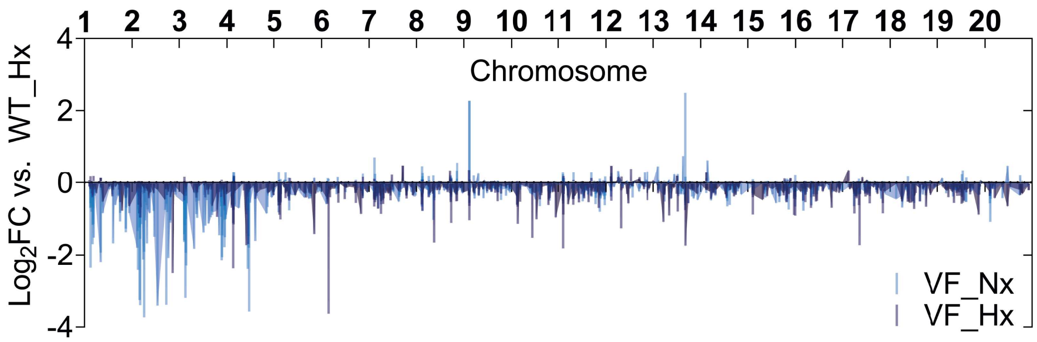


### Fig. S22

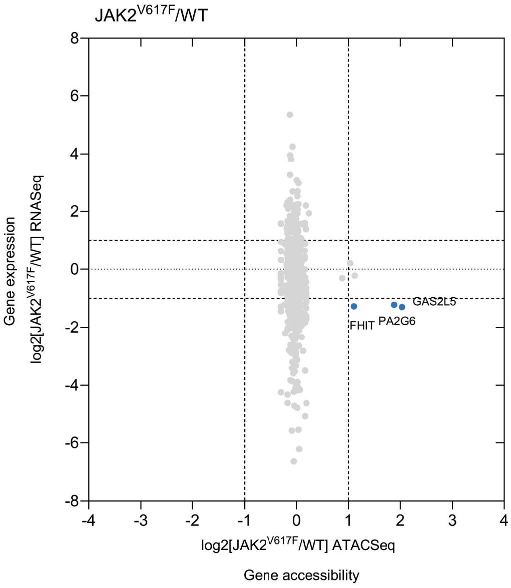


### Fig. S23

| Signature: | VF_Nx | VF_Hx |
|---|---|---|
| HIF-1 Metagene | RBPJ | RBPJ |
| | ENO1 | DDIT4 |
| | GBE1 | |
| | KDM4B | |
| | LDHA | |
| | MXI1 | |
| | PGAM1 | |
| | PGK1 | |
| | AK4 | |

### Fig. S24

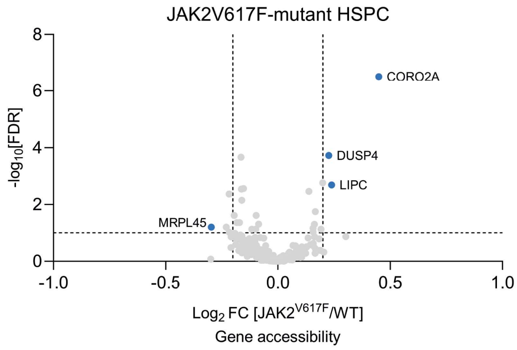

### Fig. S25

Key JAK2V617F gene signature genes

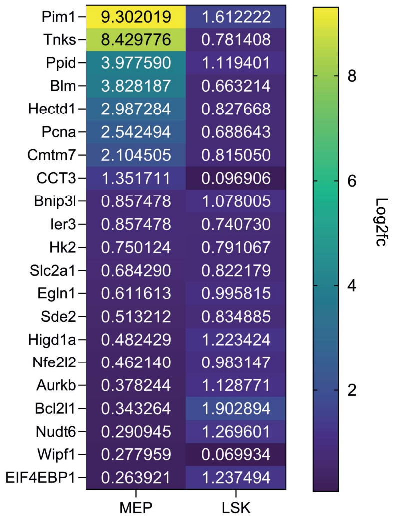


### Fig. S26

### Fig. S27

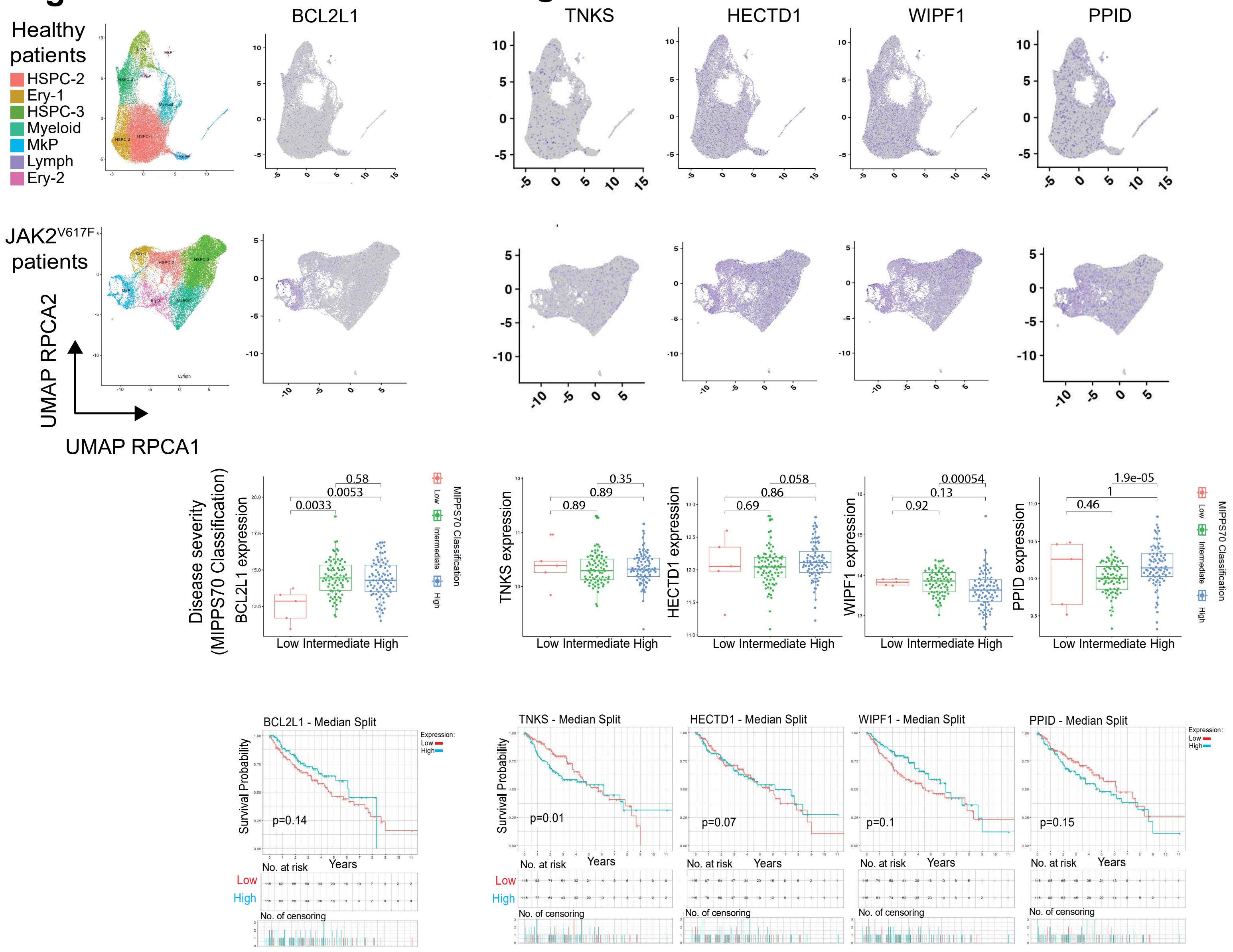

### Fig. S28

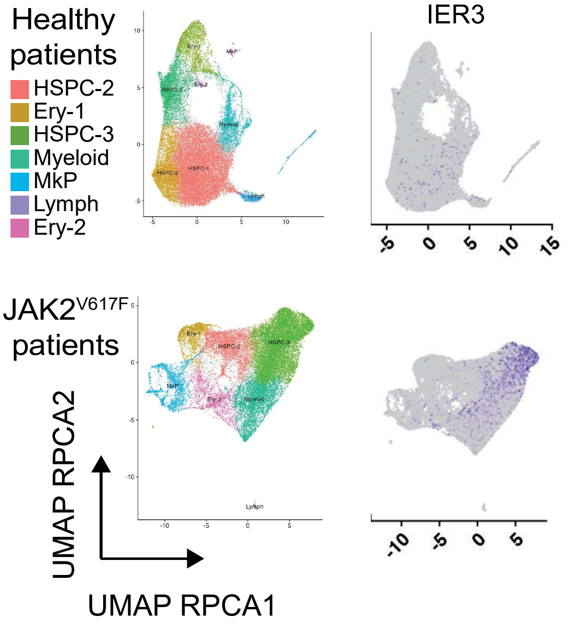


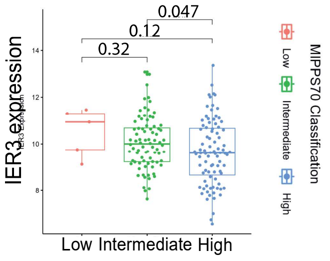


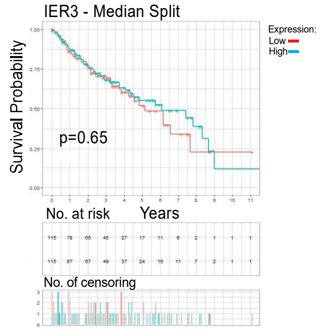


### Fig. S29

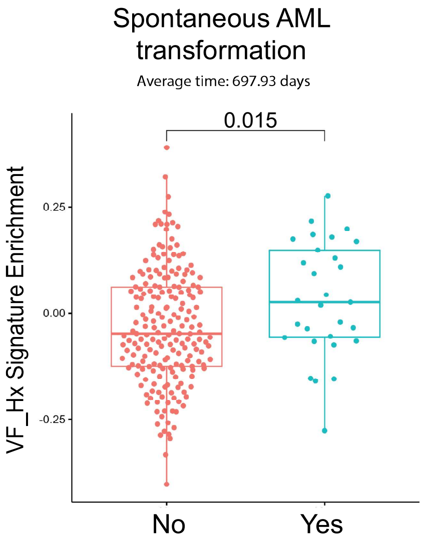


### Fig. S30

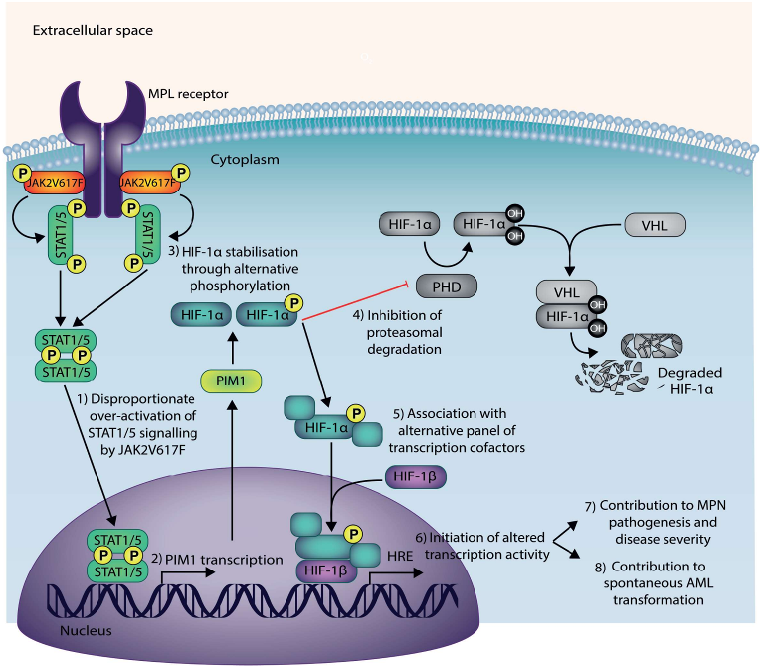
